# Comprehensive annotation and characterization of planarian tRNA and tRNA-derived fragments (tRFs)

**DOI:** 10.1101/2020.08.25.266106

**Authors:** Vairavan Lakshmanan, Dhiru Bansal, T N Sujith, Padubidri V Shivaprasad, Dasaradhi Palakodeti, Srikar Krishna

## Abstract

tRNA-derived fragments (tRFs) have recently gained a lot of scientific interest due to their diverse regulatory roles in several cellular processes. However, their function in dynamic biological process such as development and regeneration remains unexplored. Here, we identify and characterise a role for tRFs in planarian regeneration. In order to characterise planarian tRFs, we first annotated 457 tRNA loci in *S.mediterranea* combining two tRNA prediction programs. Annotation of tRNAs facilitated the identification of three main species of tRFs in planarians – the shorter tRF-5s and itRFs, and the abundantly expressed 5’-tsRNAs. Spatial profiling of tRFs in sequential transverse sections of planarians revealed diverse expression patterns of these small RNAs, including those that are enriched in the head and pharyngeal regions. Expression analysis of these tRF species revealed dynamic expression of these small RNAs over the course of regeneration suggesting an important role in planarian anterior and posterior regeneration. Finally, we show that 5’-tsRNA in planaria interact with all three SMEDWI proteins while sequence analysis revealed a possible involvement of Dicers in the processing of itRFs. In summary, our findings implicate a novel role for tRFs in planarian regeneration, highlighting their importance in regulating complex systemic processes. Our study adds to the catalogue of post-transcriptional regulatory systems in planarian, providing valuable insights on the biogenesis and the function of tRFs in neoblasts and planarian regeneration.

## Introduction

Transfer RNAs (tRNAs) canonically recognize the triplet codons on the mRNA thereby delivering appropriate amino acids to the growing polypeptide chain during protein synthesis. Emerging studies have identified tRNAs as a source for a new heterogeneous class of small RNAs called tRNA-derived fragments (tRFs). Though fragments from tRNAs were observed as early as 1970s, their physiological relevance remained largely unexplored until recently (1). In the recent years, several species of tRFs have been identified and shown to be conserved across the three domains of life. Based on the region of tRNA from which these small RNAs are processed, tRFs can be categorized as tRNA derived small RNAs (tsRNAs) or tiRNA/tRNA halves (usually 30 - 35 nts) long and other shorter (< 30 nt) fragments. There are many species of shorter tRFs such as tRF-5’s (from the 5’ arm of the tRNA), itRFs (intermediate tRFs), 3’tRFs (that correspond to the 3’ arm of the tRNA). Functionally, the different classes of tRFs regulate a multitude of cellular processes through diverse regulatory mechanism (2). tRNA halves/tiRNAs repress translation during various cellular stress responses, while tRFs function similar to miRNAs associating with *argonaute* (3–5). Most of our understanding of tRFs is in the limited context of cell culture systems. However, their role in biological processes operating at larger scales have been underexplored. Functional characterization of tRF-5s in human embryonic stem cells revealed an important role for these small RNAs in stem cell differentiation (6). Further, using heterologous models of stem vs differentiating states we previously showed that 5’- tsRNAs play an important role in regulating cell state transitions (7). tsRNAs have also shown to be abundantly expressed in sperms acting as paternal epigenetic factors thereby contributing to intergenerational inheritance (8, 9). Given these finding, tRFs are likely to have significant impact on biological processes such as development or regeneration that operate at the level of a whole organism. This study aims to place tsRNAs in the context of these processes and examine the potential roles of tsRNAs during these events.

Freshwater planarians are flatworms primarily known for their remarkable ability to regenerate any lost tissue. This ability of the planarians to regenerate is mainly attributed to the specialized of adult stem cells called neoblasts (10–12). Regeneration in planarians occurs in a sequence of cellular events such as wound closure and healing, proliferation and differentiation of neoblasts, patterning of cells to develop proportionate organs. These events are controlled by underlying gene regulatory networks that spatially and temporally control the expression of specific genes thus orchestrating a coordinated regeneration process (13, 14). Planarians possess several gene regulatory programs that are critical for neoblast function and regeneration (10, 13, 15–19). Small RNAs are one of the key regulators of gene expression. Previous studies have comprehensively characterised the expression of several microRNAs (miRNAs) during planarian regeneration (16). Knockdown of *miR-124* family of miRNAs, resulted in the mis-patterning of brain and central nervous system during regeneration (20). However, our understanding of small RNAs in planarians and regeneration are largely limited to miRNAs and piRNAs. Identification of tRFs in planarians has not been possible due to the lack of a comprehensive annotation of tRNAs.

In the present study, we annotate tRNAs in planarians combining two different tRNA prediction algorithms. Our prediction of tRNAs facilitated the identification of tRNA-derived fragment pools in planarians revealing three main tRF species in planarians - 5’tRFs, itRFs and 5’-tsRNAs. Bioinformatic analysis of these small RNA from sequential transverse sections uncovered diverse spatial expression patterns for these small RNAs across the planarian body indicating body-wide functional relevance. Further, analysis of the previously published small RNA dataset during planarian anterior and posterior regeneration, identified tRFs enriched during various stages of regeneration highlighting a crucial role for these small RNAs in regulating the various events during regeneration. Lastly, using the existing SMEDWI pulldown data, we observed 5’-tsRNAs interact with all the SMEDWI proteins in planaria, thus offering a possible biogenesis and functionality for these small RNAs. Further, sequence analysis of itRFs revealed an enrichment for ‘U’ at the first base hinting at Dicer-based cleavage of these small RNAs. Our study, for the first time, identifies a novel class of small RNAs in planarians thus expanding the post-transcriptional regulatory systems that govern stem cell function and regeneration.

## Results

### Annotation of tRNAs and codon usage in planarians

To predict tRNA genes in *Schmidtea mediterranea*, we used tRNA prediction algorithms tRNAScan-SE and Aragorn to scan dd_Smes_G4 (21) version of genome. tRNAScan-SE predicted 4,115 putative loci while Aragorn predicted 4,143 loci across planarian genome (Figure 1A). Sequences predicted by both the programs were clustered and overlapped using CD-HIT (>90% sequence similarity) (22). Clustering these predictions narrowed down the tRNAs to 708 unique sequences (Figure 1A). Further filtering these 708 sequences using tRNAScan-SE (v2.0), an improvised predictive algorithm and classifier, resulted in the prediction of 457 unique tRNA genes in planrians (23). The identified 457 tRNA genes were classified into three categories (Figure 1A), (i) Standard tRNA genes, which code for the standard 20 amino acids, (ii) UNDET tRNAs, that are potential tRNA genes for which the programs were unable to assign a triplet codon confidently. (iii) Pseudo tRNA genes; that are derived from tRNAs with point mutations, insertions or deletions. The UNDET tRNAs predicted in our analysis were resolved further using TFAM (24) to assign the amino acids these tRNAs could code for based on sequence conservation with known tRNAs across all organisms. These 457 tRNA genes were represented across 1372 loci on the planarian genome, with Lysine having highest number of tRNA genomic loci (104 loci for 32 tRNA genes, S1A). The predicted planarian tRNAs had a median length of 75 nts with less variation across the three categories of tRNAs (Figure 1B) comparable to those observed across eukaryotes (with a median length around 72/73bp) (Figure S1B). Analysis of the anticodon positions revealed that ~30% of planarian tRNA anticodons originate at the 34^th^ nucleotide on the tRNA (Figure 1C). Sequence analysis of the 457 predicted planarian tRNA sequences revealed high sequence conservation between the tRNAs carrying same amino acids (Extended Supplementary 1)

**Figure 1:**
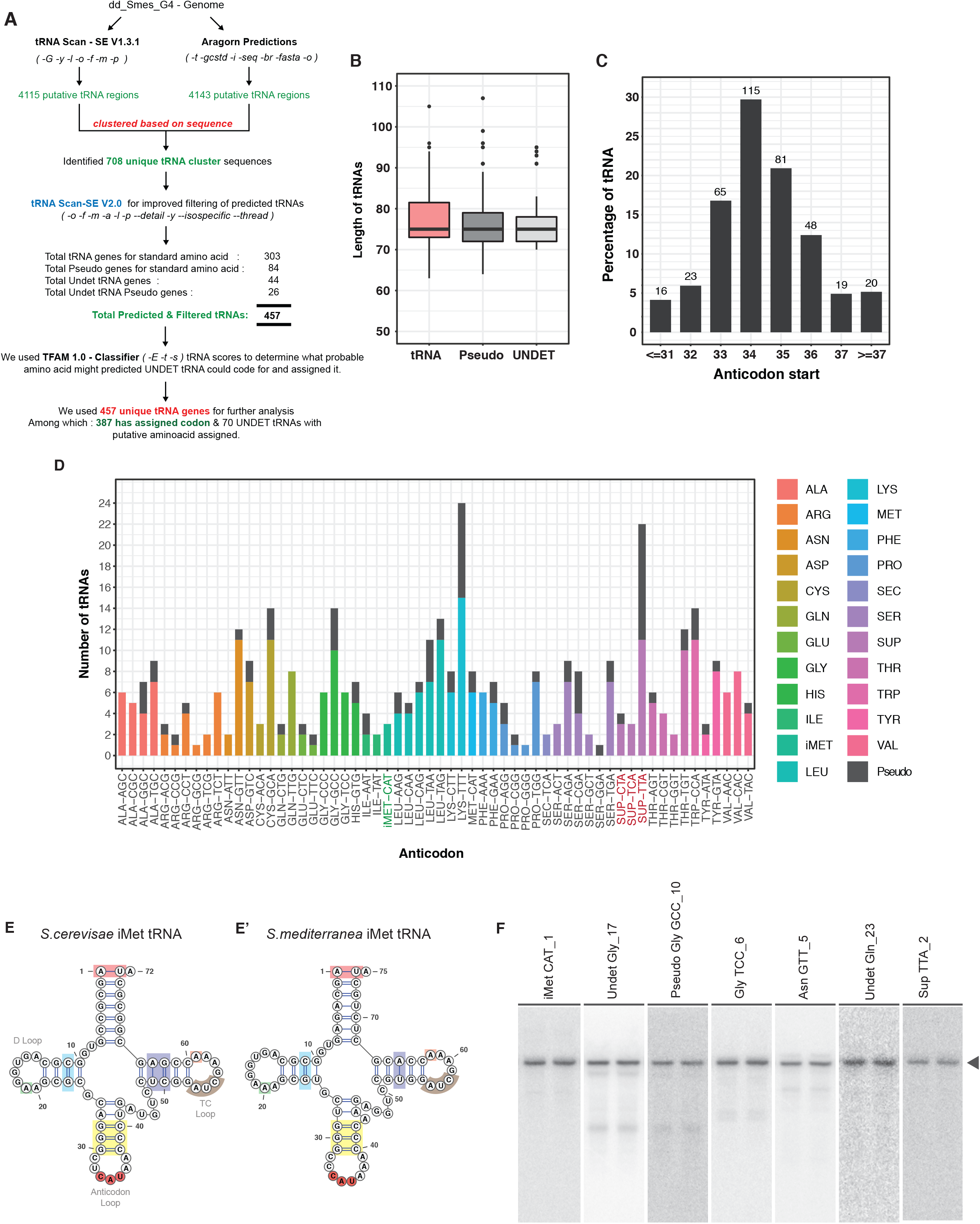
tRNA annotations in planaria. A) Bioinformatic pipeline used to annotate planarian tRNAs. Two different prediction algorithms were used to identify a confident set of 457 tRNAs in *S.mediterranea*. B) Median lengths of tRNA identified in *S.mediterranea*. C) Bar graph depicting the position of anticodon across planarian tRNAs. D) The gene copy numbers of all the identified planarian tRNAs across *S.mediterranea* genome. E) Conserved features if i-Met tRNAs identified in planaria as compared to the *S.cerevisae* iMet tRNA. The colours indicate the conserved sequence signatures. F) Northern hybridization blot validation of predicted tRNAs (shown in duplicates). The arrow points to the tRNA band.

Among the 457 tRNAs, 347 sequences code for standard 20 amino acids (including UNDET tRNAs) and 110 sequences were classified as pseudo genes (Figure 1D, S1A, S1C and Table S1). The total number of tRNA genes predicted is comparable with well annotated species (human, mouse, chick, rat and fly) (Figure S2A). From our stringent prediction, we identified tRNA genes carrying anticodons against 54 standard codons out of the 61 total codons (Figure 1D and S1C). Among the tRNAs coding for methionine, we were able to identify three initiator methionine tRNAs (tRNA iMet-CAT) in the *S.mediterranea* genome, based on certain conserved sequence features that exists across all the eukaryotic iMet-tRNAs (Figure 1E and E’). Further, we also identified selenocysteine and three suppressor tRNAs (SUP-CTA, SUP-TCA and SUP_TTA) in *S.mediterranea*. Suppressor tRNAs are class of tRNAs that can recognize stop codon (25, 26). This class of tRNAs alleviate pre-mature termination of protein synthesis due to mutations in standard tRNA anticodon that results in a stop-codon (25, 26). To validate our prediction of tRNAs, we performed Northern hybridizations for candidate tRNAs belonging to the different tRNA categories (iMet-CAT, UNDET-Gly_17, UNDET-Gln_23, Pseudo Gly-GCC, Asn-GTT, and SUP-TTA_3). Northern blots revealed the expression of all the 5 tested tRNAs evidenced by a prominent tRNA band (Figure 1F). Collectively, our prediction pipeline identified 457 tRNAs genes in the planarian genome, which exhibited sequence signatures and characteristics conserved across organisms.

Studies across metazoans have shown that the 64 codons are used at different capacities and these preferences vary across different biological contexts (27–31). One of the evolutionary determinants of codon use, and thereby translational efficiency, in organism is the abundance of tRNAs (32). Since determining the tRNA abundance has been difficult owing to the high degree of modifications on tRNA, efforts to understand tRNA-codon relations have used tRNA gene copy numbers as a proxy (33–35). In order to study the relation between the identified tRNA genes and the codon usage in *S.mediterranea*, we first calculated the codon usage using recently annotated transcriptome (36). Codon frequency analysis revealed Lys-AAA, Asn-AAT and Glu-GAA to be the most used triplet codon across the planarian transcriptome (>50% codon frequency per 1000 codons) (Figure S2B and Table S2). In agreement, Lys-TTT tRNA(the cognate pair for AAA codon) exhibited the highest tRNA gene copy number, while other highly gene copy number tRNAs (such as Asn-GTT, Gly-GCC and Val-CAC) showed inverse correlations with its cognate codon usage frequency (Figure S2B). However, correlation between the planarian isoaccepting-tRNA gene copy number and the amino acid usage revealed a positive linear relationship with a correlation of 0.54, comparable to those observed across all three domains of life (Figure S2C) (35). Probing these dynamics might shed more light in understanding mechanism that regulates translation in planarians.

### tRNA-derived fragments (tRFs) in Planarians

Using our annotated planarian tRNAs, we sought to identify the tRNA-derived fragments in planarians. To obtain a holistic understanding of these small RNA in planarians, we analysed our previously published planarian small RNA data (16). Initially, reads from planarians (intact whole animal) were mapped to our annotated tRNA sequences. Although miRNAs and piRNAs were the most dominant small RNA species (20.6 and 17.2 % respectively), 2.04% of reads (0.21 million reads of the 10.3 million total reads mapping to genome) mapped to our newly annotated tRNA database (Table S3). Encouraged by these observations and in order to mine tRFs with greater sequencing depth in planarians, we sectioned planarians into 12 pieces (11 sequential sections from head to tail with pharynx as the 12^th^ part), a procedure termed as salami sectioning (37). Deep sequencing of small RNAs from each of these individual sections was performed. Comparison of the tRFs from the whole animal to the averaged reads of the salami section (a proxy for the whole animal) revealed a high correlation (R^2^= 0.93) highlighting the robustness of our dataset (Figure S3A). Subsequently, we mapped 18-35 nt reads obtained from salami sections to miRNAs, piRNAs, tRNAs etc. (Figure 2A). The overall small RNA (18-35nt) reads that mapped to our annotated tRNAs ranged from 0.8 to 3.91% across the salami sections (Table S3). Though the reads mapping to tRNAs represent only a small fraction, their individual expression levels were comparable to some of the highly expressed planarian miRNAs (Figure S3B).

**Figure 2:**
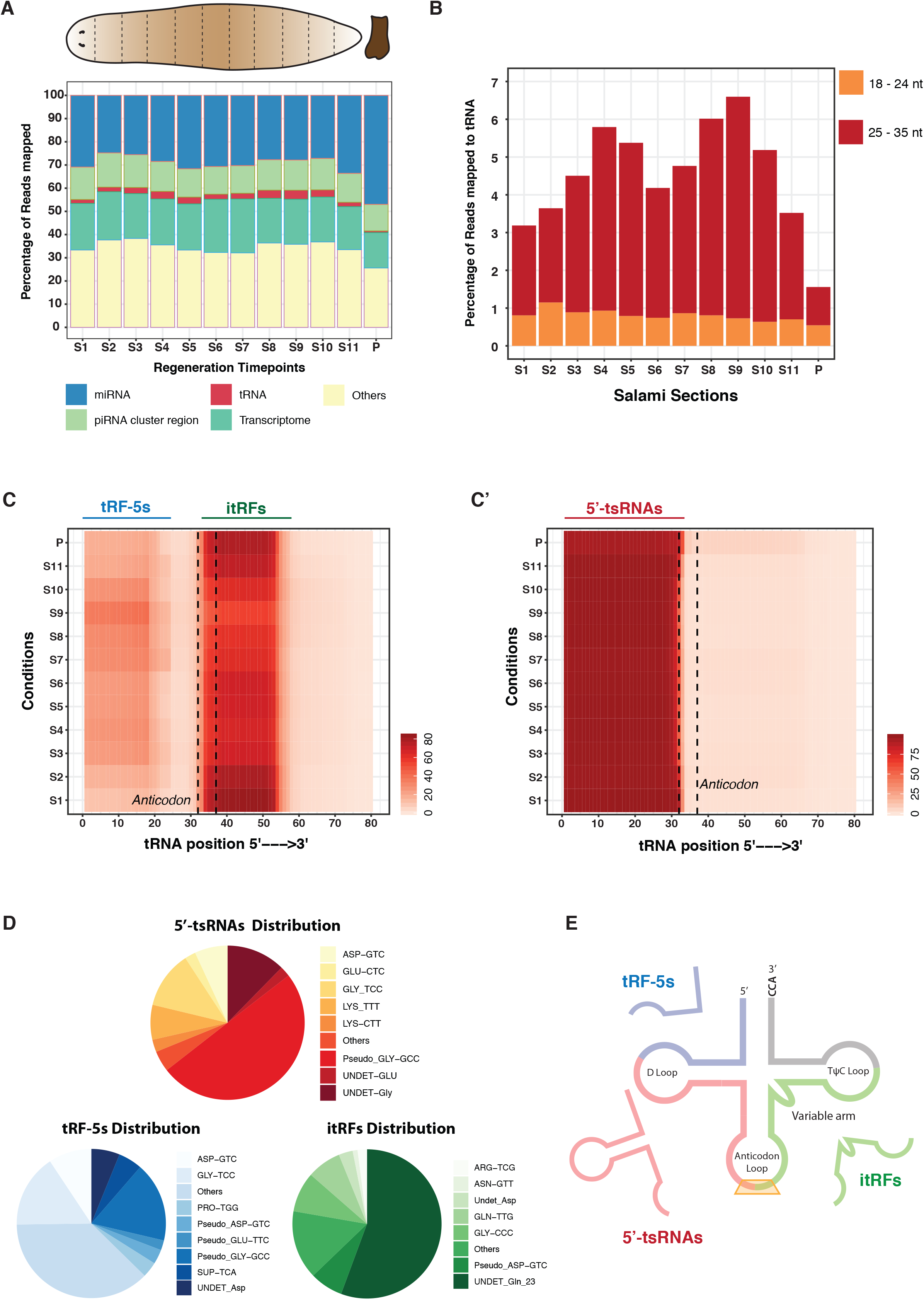
Identification of tRNA-derived fragments across planarian body axis. A) Percentage distribution of all the identified small RNAs across the sections of planarian body. B) Percentage distribution of small RNA reads, mapping to tRNAs across planarian body sections. Small RNA reads are segregated into 18-24 nts and 25-35 nts. C) Perbase coverage of 18-24 nt reads across parent tRNA length. C’) Perbase coverage of 25-35 nt reads across parent tRNA length. D) Piechart depicting the percentage disctribution of 5’-tsRNAs, tRF-5s and itRFs in planaria. E) Pictorial depiction of the three tRNA-derived fragments identified in this study.

Previous studies in other organisms have classified tRFs based on the length and the region of the parent tRNA from which these fragments originate (2). We grouped small RNA reads that mapped to planarian tRNAs based on size as 18-24 nts and 25-35 nts. Among the two fragment size distributions, we observed 25-35 nt to be the dominant population with ~1 to 5.8% reads mapping to tRNAs as compared to the 0.5 to 1.1% of 18-24nt reads (Figure 2B and Table S3). Further, to understand the region on the tRNA from which these small RNAs arise, we plotted the per base coverage for these two sized population over the total length of the parent tRNA. Perbase coverage of the 18-24 nt species showed enrichments for two different tRNA fragment species; the tRF-5s, that originate from the 5’ end of the tRNA; and the itRFs, those that originate from the anticodon region and extend into the 3’ arm of the tRNA (Figure 2C). Moreover, per base coverage for the 25-35 nts reads across the length of the tRNA suggested these reads almost exclusively arise from the 5’ half of the tRNA. Our analysis also revealed that among the 18-24 nt species of itRFs and tRF-5s, the itRFs are processed as a homogenous size of 20 nts whereas tRF-5s were processed into three dominant size pools - 18 nt, 21 nt and 24 nts (Figure S3C). Further analysis of these three different species of tRNA fragments revealed that the majority of reads for the tRF-5s, itRFs and 5’-tsRNAs correspond to a specific set of tRNAs, an observation made in other organisms as well (7, 8, 38). Majority of the 5’-tsRNAs were processed from tRNA Pseudo-GlyGCC contributing to ~50% of the total 5’-tsRNA (Figure 2D and Table S4). Northern hybridizations validated the expression of the top two abundantly expressed 5’-tsRNAs, Pseudo-GlyGCC and UNDET-Gly (Figure S3D). Similarly, among the itRFS, reads mapping to tRNA UNDET-Gln23 contributed to ~56% of the reads. However, reads were evenly distributed among the top tRNAs that generate tRF-5s (Figure 2D).

Based on the tRNA to which the reads mapped, we observed that 117 planarian tRNAs are processed into all the three fragments with at least one read mapping to the parent tRNA (Figure S3E). Further filtering these tRNAs based stringent read cut off we identified 12 tRNAs that could be processed into all three fragments (compared to 117 without cut-offs). 25 tRNAs are capable of producing the both tRF-5s and the 5’-tsRNAs species while 11 tRNAs are capable of producing itRFs and 5’-tsRNAs (Figure S3E’). Interestingly we observed that specific groups of tRNAs are uniquely processed into a particular species of tRNA fragment; 47 tRNAs produce only itRF species, while 41 tRNA produce tRF-5s and 3 tRNAs are processed into only the 5’-tsRNA species (Figure S4F’ and Table S5). It was previously reported that the the abundance of tRNAs dictate the utilization of tRNA to generate tRFs (39). Notably in planaria, some of the highest expressed tRFs, were processed from specific isodecoder-tRNAs that displayed high gene copy numbers (Asp-GTC and Asn-GTT) with a corresponding low codon usage across the planarian genome (Figure S2B). Together, our analysis of tRNA-derived fragments in planarians identifies three different species of tRFs; the smaller tRF-5s, itRFs and the more abundant and longer 5’-tsRNAs (Figure 2E).

### Spatial expression patterns of tRFs across anterior-posterior (AP) axis of S.mediterranea

We next profiled the expression of small RNAs across planarian salami sections. Initially, we analysed the expression profiles of the most studied class of small RNAs, the miRNAs. Our analysis identified four distinct spatial clusters for miRNAs (Figure S4A). Consistent with previous reports, our salami section strategy showed enrichments for miR-124, a brain enriched-miRNA, in the anterior sections of planarians (Figure S4B). Subsequently, we studied the expression of the identified tRFs across the AP axis of planarians. The expression of the dominant 5’-tsRNA species could be broadly clustered into 4 domains. The first cluster of 5’-tsRNAs (Cluster-1), such as Gly-TCC_2, Lys-TTT_8 and Lys-TTT_15, showed enrichments in pre- and post-pharyngeal regions with low expression in the head, tail and the pharynx (Figure 3A and Table S4). The expression of 5’- tsRNAs belonging to the second cluster exhibited profiles similar to the expression of neoblast-specific transcripts, such as *smedwi-1, smedwi-2, smedwi-3, vasa* and *bruli* (Figure S4C). The third cluster of 5’-tsRNAs were expressed uniformly across the different sections with higher expression in the pharynx (P). Some of the examples of pharynx enriched 5’-tsRNAs are Undet-Gln_23, Gln-TTG_2, Thr-AGT_1 etc. (Figure 3A). This cluster of 5’- tsRNAs (Cluster-3) were grouped based on their high expression in the head region (S1) suggesting that these could be expressed in the tissues of the head such as brain, eyes etc. (Figure 3A). Interestingly, although these tsRNAs were enriched in the head, their expression domains sometimes extended beyond the head regions leading us to speculate that these tsRNAs may be expressed in the CNS or specific neurons. The fourth cluster of 5’- tsRNAs varied in expression across the sections with no definitive region of high expression. Similarly, clustering tRF-5s and itRFs based on expression profiles across salami sections resulted in three broad clusters alike 5’- tsRNA clusters – those that are enriched in the head region, the pharynx region or in certain sections apart from these two regions (Figure 3B and 3C and Table S4). Our expression analysis of tRFs thus identifies distinct spatial expression of these small RNAs suggesting they may be important for varied systemic functions in planarians.

**Figure 3:**
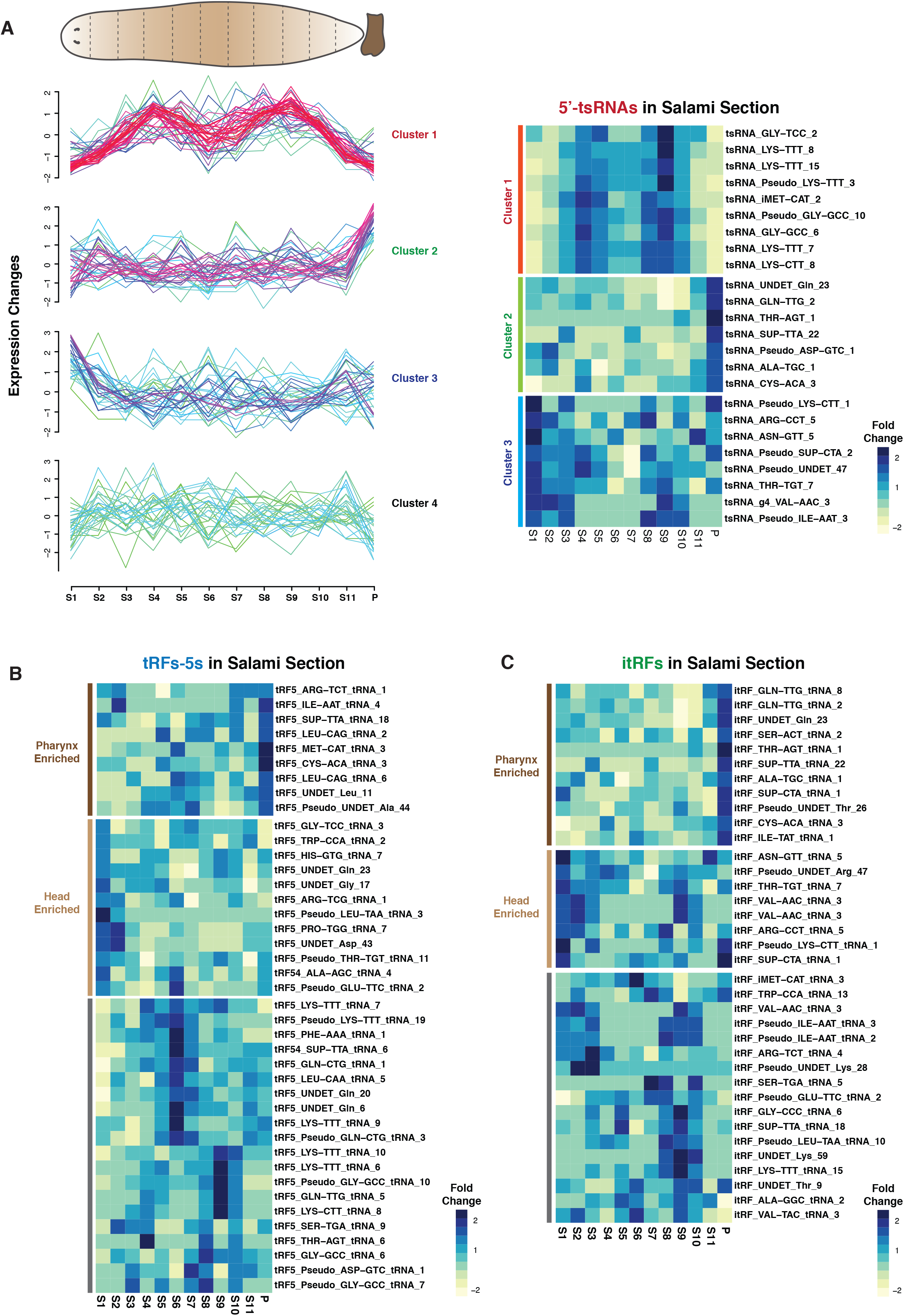
Spatial profiling of tRNA-derived small RNAs across planarian salami sections. A) Expression profiles of 5’-tsRNAs across the different sections of planarian body identifies three clusters of expression. In soft clustering, membership value represents how well a gene is represented in a cluster. The red and purple color represents candidates with high membership value while yellow or green indicates candidates with low membership value. A) Heatmaps of candidate 5’tsRNAs that show various patterns of observed expression. B) Heatmaps of candidate tRF-5s across salami sections. The expressions are grouped based on high expression in head region, pharynx or orther regions of the planarian body. C) Heatmaps of candidate itRFs across salami sections. The expressions are grouped based on high expression in head region, pharynx or other regions of the planarian body.

### Expression of 5’-tsRNA during Planarian regeneration

In our previously published work on small RNAs in planarian regeneration, we identified clusters of miRNA expression in the regenerating tissue (blastema) across various time points (3 hrs, 6 hrs, 12 hrs, 1 day, 3 day, 5 day and 7 day) of anterior and posterior regeneration (Figure 4A). This dataset was used to analyze the expression of different tRNA fragment species in planarians. As observed in salami sections, we obtained 2 to 5.4% of total reads (18–35) mapping to tRNAs (Figure 4B and Table S7). Preliminary analysis revealed dynamic changes in the different species of tRFs in both anterior and posterior regeneration. Over the course of anterior and posterior regeneration, the overall population of tRFs seem to increase in 3 hours post amputation (hpa) and reverts back to uncut levels over the 7-day regeneration regime (Figure 4C). Similar to the size distributions of tRFs observed across salami sections, we found that 25-35 nt reads were the most abundant species during regeneration (Figure S5A). Per base coverage of reads that map to tRNA revealed that all the three species; itRFs, tRF-5s and 5’- tsRNAs are also expressed during regeneration (Fig. S5B and S5B’). Further, the overall expression pattern of these three species, revealed a divergent expression at early hours post amputation (Figure 4D). While the collective 5’-tsRNA population doubled at 3hrs post amputation, the tRF-5 species remained unchanged over this time points, and the itRFs levels decreased two-fold in both paradigms of regeneration. This divergent expression for the three species of tRFs suggests distinct functionalities for these populations during regeneration.

**Figure 4:**
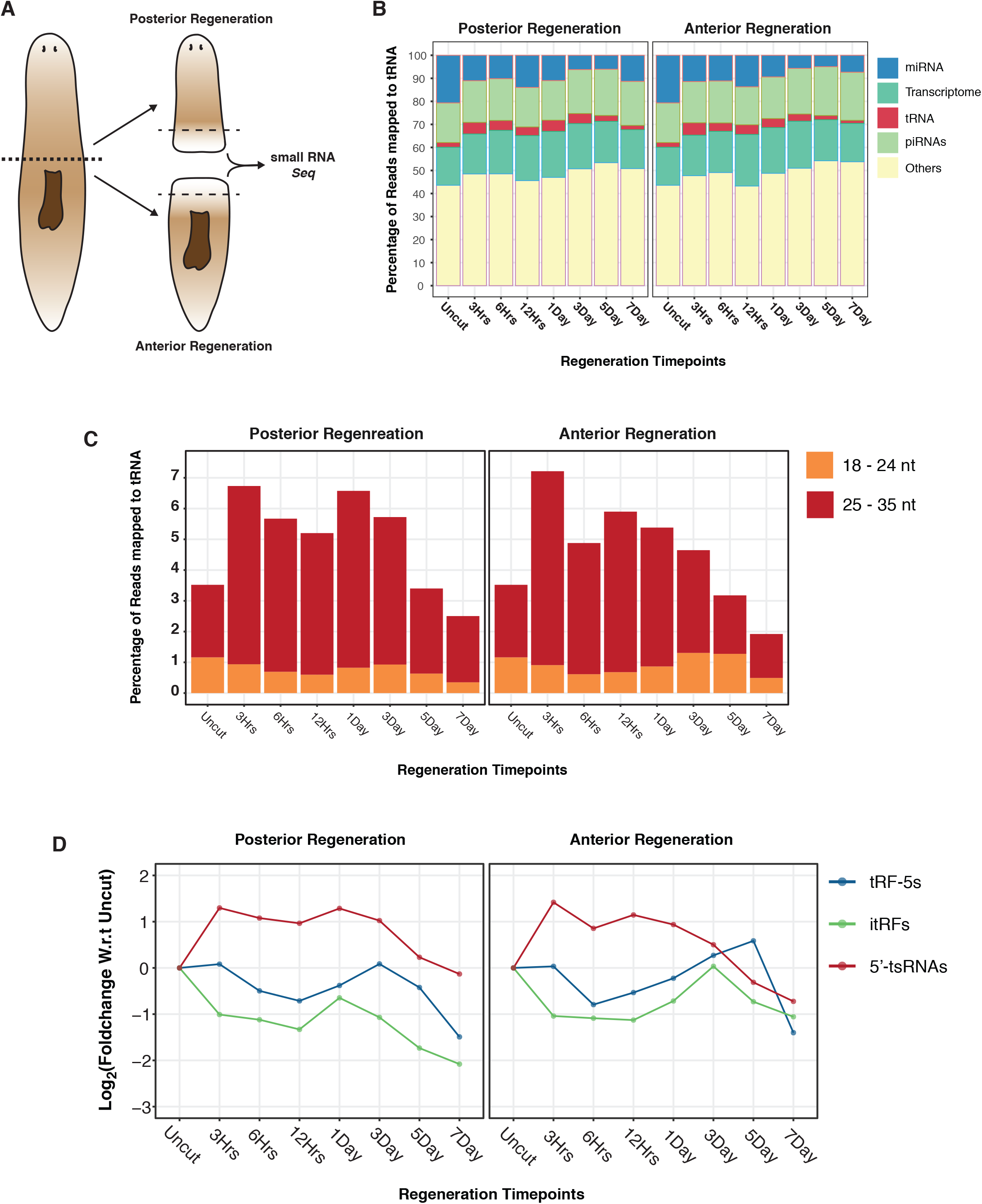
tRNA-derived fragments in planarian anterior and posterior regeneration. A) Schematic describing the strategy for sequencing. B) Percentage distribution of all small RNA populations across various timepoints of anterior and posterior regeneration. C) Percentage distribution of small RNA reads, mapping to tRNAs across planarian anterior and posterior regeneration. Small RNA reads were segregated into 18-24 nts and 25-35 nts. D) Expression profiles of each tRNA-derived fragment population over timepoints of anterior and posterior regeneration shows divergent patterns.

We next investigated the expression of individual 5’-tsRNAs, tRF-5s and itRFs over the course of regeneration (Table S8). Analysis of 5’-tsRNA expression across regenerating time points showed distinct clusters. The clusters were demarcated as Anterior Regeneration Cluster (ARC) and Posterior Regeneration Cluster (PRC). During posterior regeneration our analysis revealed that 5’-tsRNAs followed 4 main expression profiles (Figure 5A). PRC-1: represents the 5’-tsRNAs that are upregulated in early time points of regeneration such as 3 hpa and their expression gradually decreases over later time points of regeneration (3 dpa – 7 dpa). 5’- tsRNAs belonging to PRC-2 showed decreased expression in early time points of regeneration and peaked around 12 hpa (Figure 5A). The expression of this cluster of 5’-tsRNAs correlates with the ‘second wave of wound healing response’ as reported by Wenemoser *et al.,* suggesting that these 5’-tsRNAs could be involved in the wound healing and early regeneration response. PRC-3 comprises of 5’-tsRNAs that exhibit either decreased or no change in the expression till 3 dpa beyond which their expression increases (Fig. 5A). These PRC-3 5’-tsRNAs may be involved in the later stages of development and differentiation programs that set in 3 dpa during regeneration. Lastly, we identified a unique cluster (PRC-4) of 5’tsRNAs that are specifically downregulated at 12 hpa but are comparable to uncut levels at other timepoints of regeneration. This cluster of 5’-tsRNAs could possibly mediate the transitions between early and late regeneration programs. We were also able to classify a unique set of 5’-tsRNAs, based on their temporal expression, as “early” (3hrs-1day) regeneration and “late” regeneration (3days – 7days) 5’-tsRNAs (Figure S6A and S6A’).

**Figure 5:**
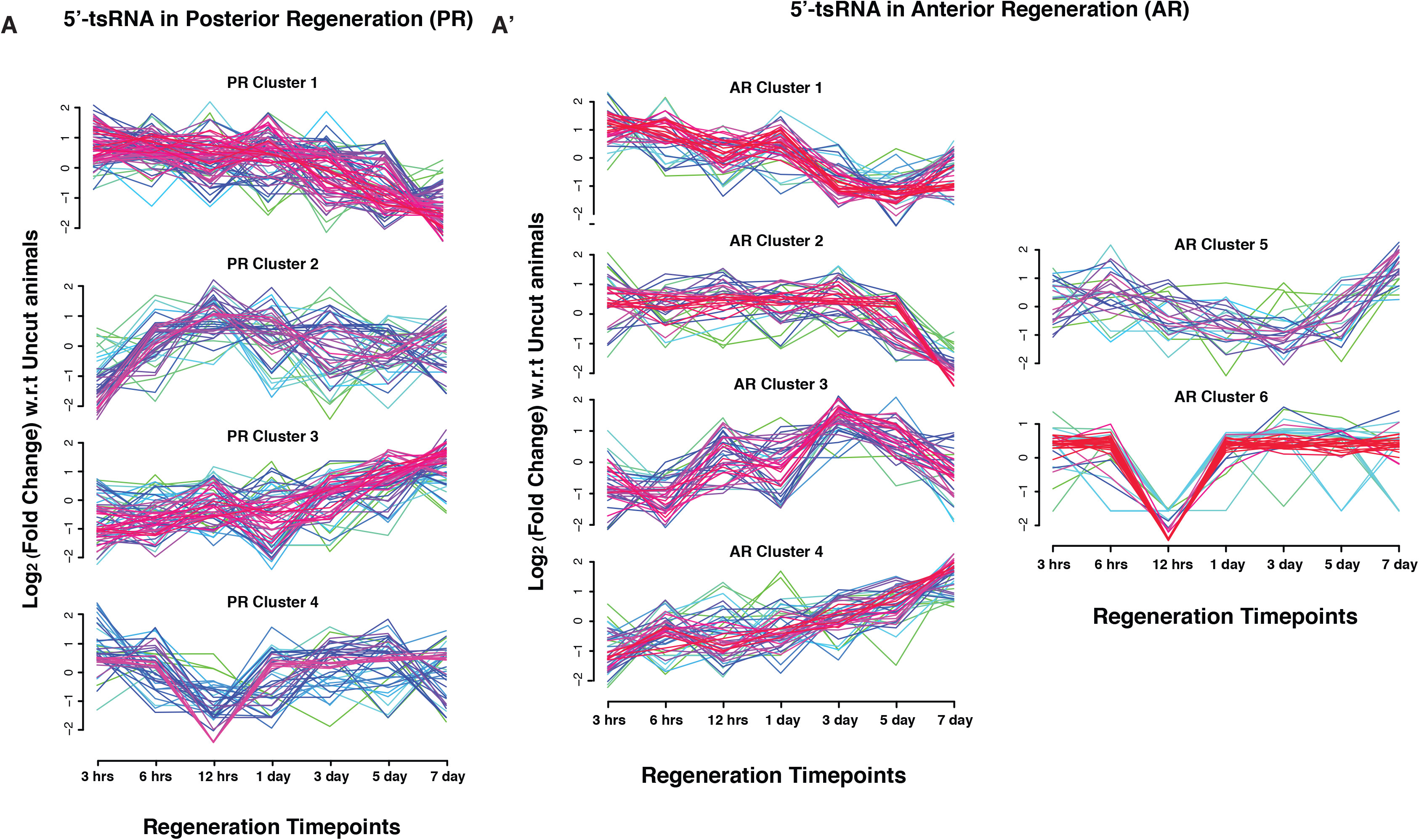
Expression profiles of 5’-tsRNAs across planarian anterior and posterior regeneration. A) Expression profiles of 5’-tsRNAs across the different timepoints of posterior regeneration clustered into similar expression types. A’) Expression profiles of 5’-tsRNAs across the different timepoints of anterior regenerating fragment clustered into similar expression types.

We also examined the expression of 5’-tsRNAs in anterior regenerating animals. As compared to the posterior regeneration tsRNAs, the expression of the 5’-tsRNAs during anterior regeneration could be confidently segregated into 6 clusters of expression (Figure 5A’). The ARC-1 and −2: comprises of 5’-tsRNAs that has enhanced expression during early time points of regeneration (3hrs onwards). ARC-1 cluster showed gradual decrease in the expression of 5’-tsRNAs, plummeting subsequently after day 1 post amputation (Figure 5A’). ARC-2 5’-tsRNAs have a more prolonged period of elevated expression with a sharp decrease in expression at 7 dpa. The trends exhibited by ARC-1 and ARC-2 classes of 5’-tsRNAs would suggest a role for these small RNAs in regulating the early events of regeneration. ARC-3 and ARC-4 displayed similar expression profiles both showing increased expression patterns at later time points of regeneration. Both these clusters showed decreased 5’-tsRNA expression in early time points of regeneration with a gradual or a sharp increase in expression around 1-3 dpa (Figure 5A’). While ARC-3 includes 5’-tsRNAs that peak in expression around 3-5 dpa followed by a decrease at 7 dpa; the ARC-4 showed highest expression at 7 dpa similar to the expression of the PRC-3. We speculate that ARC-3 clusters could represent the wave of 5’-tsRNAs that are essential for the formation of the head structures while the ARC-4 cluster could represent the pool that could be involved in homeostasis, growth and organization of the head structures. ARC-5 displayed increased expression at early (3hpa and 6 hpa) and late (7 dpa) however they displayed decreased expression at 3-5 dpa (Figure 5A’). Here, we speculate that ARC-3, ARC-4 and ARC-5 could be the 5’-tsRNAs that are involved in later stages of head regeneration. We also identified a set of 5’-tsRNAs (ARC-6) that showed a sharp reduction in expression at the 12 hrs time point akin to the PRC-4 (Figure 5A’).

Next, we profiled the expression patterns of tRF-5s and itRFs at different time points of anterior and posterior regeneration. Similar to the expression patterns observed with 5’-tsRNAs, we were able to categorize the expression of tRF-5s and itRFs into early and late waves of expression (Figure 6A, 6A’, 6B and 6B’ and Table S8). Our analysis also revealed a group of 5’-tsRNAs potentially not involved in the regeneration process as evidenced by the downregulation of these small RNA over all the tested regeneration time points (Figure 6A, 5A’, 6B and 6B’). To distinguish the tRFs that are specific to either anterior or posterior regeneration, and those that are involved in a common regenerative program (irrespective of anterior or posterior regeneration), we analyzed the overlapping tRFs between these two paradigms. Interestingly, during the early regeneration response, a greater percentage of tRFs were common between anterior and posterior regeneration (around 25% for itRFs, 50% for tRF-5s and 45% for 5’-tsRNAs) (Figure S6B). However, during the late response, a large percentage of tRFs emerged specific to head regeneration, possibly those that are important for the regeneration of the head structures. We also observed a concomitant decrease in the common tRFs (for tRF-5s and 5’-tsRNAs) and tRFs specific to the tail regeneration (Figure S6B). Collectively, our analysis identified dynamic expressions of several tRFs throughout planarian regeneration.

**Figure 6:**
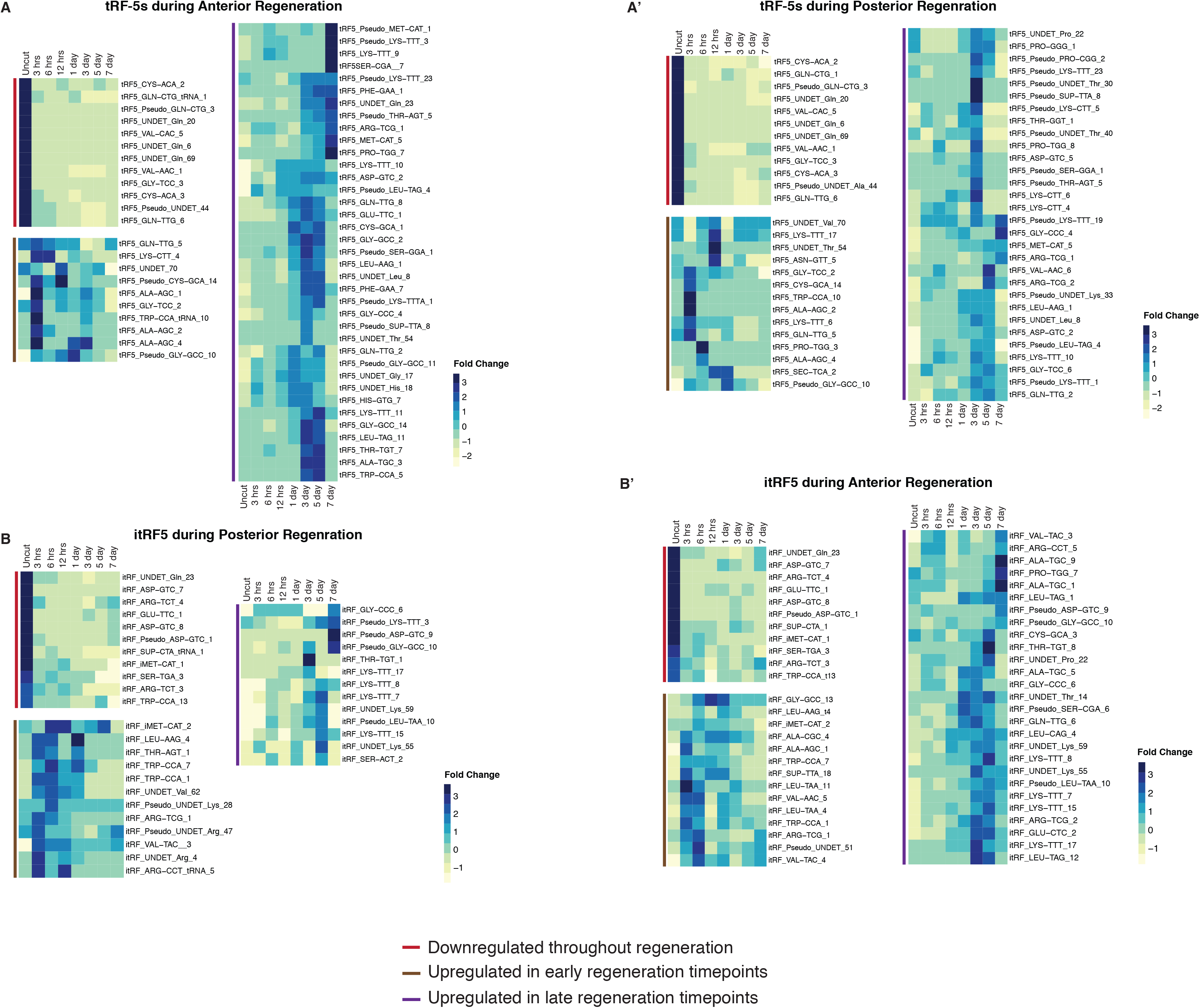
Expression profiles of tRF-5s and itRFs across planarian anterior and posterior regeneration. A and A’) Heat maps of candidate tRF-5s during anterior and posterior regeneration. We identified three groups of expressions. tRF-5s that are downregulated over all regeneration timepoints, tRF-5s that are upregulated in early timpoints of regeneration (3rs −1day) and late timepoints of regeneration (3days - 7days). B and B’) Heat maps of candidate itRFs during anterior and posterior regeneration. We identified three groups of expressions. itRFs that are downregulated over all regeneration timepoints, itRFs that are upregulated in early timpoints of regeneration (3rs −1day) and late timepoints of regeneration (3days - 7days).

### Planarian 5’-tsRNAs interact with all three SMEDWIs

Several studies have identified potential enzymes and factors responsible for processing of tRFs (3, 6, 40). Of the different species of tRFs, processing of tRNAs by angiogenin to produce tsRNAs (or tiRNAs or tRNA halves) under stress conditions has been the most studied (3, 41). Sequence homology based survey suggested that *S.mediterranea* lacks proteins homologous to angiogenin. This implied that the tsRNAs in planarians are processed by a completely different mechanism. Recent evidences in mammalian system have also arrived at similar conclusions that 5’-tsRNAs could be processed in an angiogenin-independent manner (7, 42). Another protein that is abundantly expressed in planarians and one that has been reported to associate with 5’-tsRNAs in other systems is Piwi (43). Planarians express three piwi proteins SMEDWI-1, SMEDWI-2 and SMEDWI-3 (11, 12, 44). A recent study in planarians identified the RNAs associated with the three SMEDWI proteins to understand their function (44). We mapped this dataset to our annotated tRNAs to explore if 5’-tsRNAs associate with these proteins. The reads that mapped tRNAs were predominantly of the size 30-35 nts suggesting that SMEDWIs interact with tsRNAs (Figure S7A). Our analysis revealed that a small subset of PIWI-interacting RNAs map to tRNAs with SMEDWI-2 showing the most association (Figure 7A and Table S9). It is not surprising that tsRNAs represent only a minor fraction of PIWI-interacting RNAs considering the fact that tsRNAs make-up a small fraction of the total small RNAs in planarians, and piRNAs being the primary interactors of PIWI proteins. However, it is important to note that 80-90% of all the planarian 5’-tsRNAs associate with the SMEDWI proteins indicating PIWIs to be essential for the biogenesis and/or functioning of 5’-tsRNAs in planarians (Figure 7B). Our analysis also revealed that all the SMEDWI proteins in planarians associate with a large pool of common 5’-tsRNAs (Figure 7C). This observation suggests two possibilities, that 5’-tsRNAs employ different routes of biogenesis/function through PIWI proteins, and planarian 5’-tsRNAs may interact differentially with these three proteins across the three planarian cell populations. To explore the latter, we investigated if there is any correlation between the 5’-tsRNAs that associate with the three PIWI proteins and the tsRNAs expressed in the three different cell populations in planarians (X1, neoblasts; X2, largely comprising of progenitors and Xins the differentiated cells). To perform this analysis, we used our previously published small RNA data data from X1, X2 and Xins cell populations (16). Our analysis revealed that 5’-tsRNAs associated with SMEDWI-3 showed the highest level of correlation with these three cell types (Fig. 7C and Table S10). It has been previously reported that SMEDWI-3 is also expressed in differentiated cells suggesting the reason for stronger correlation with X2 and Xins populations compared SMEDWI-1 and 2 (44, 45). SMEDWI-1 associating 5’-tsRNAs correlated the least with the three cell types. In conclusion, our data suggests that SMEDWI proteins could be critical for planarian tsRNA biogenesis and function.

**Figure 7:**
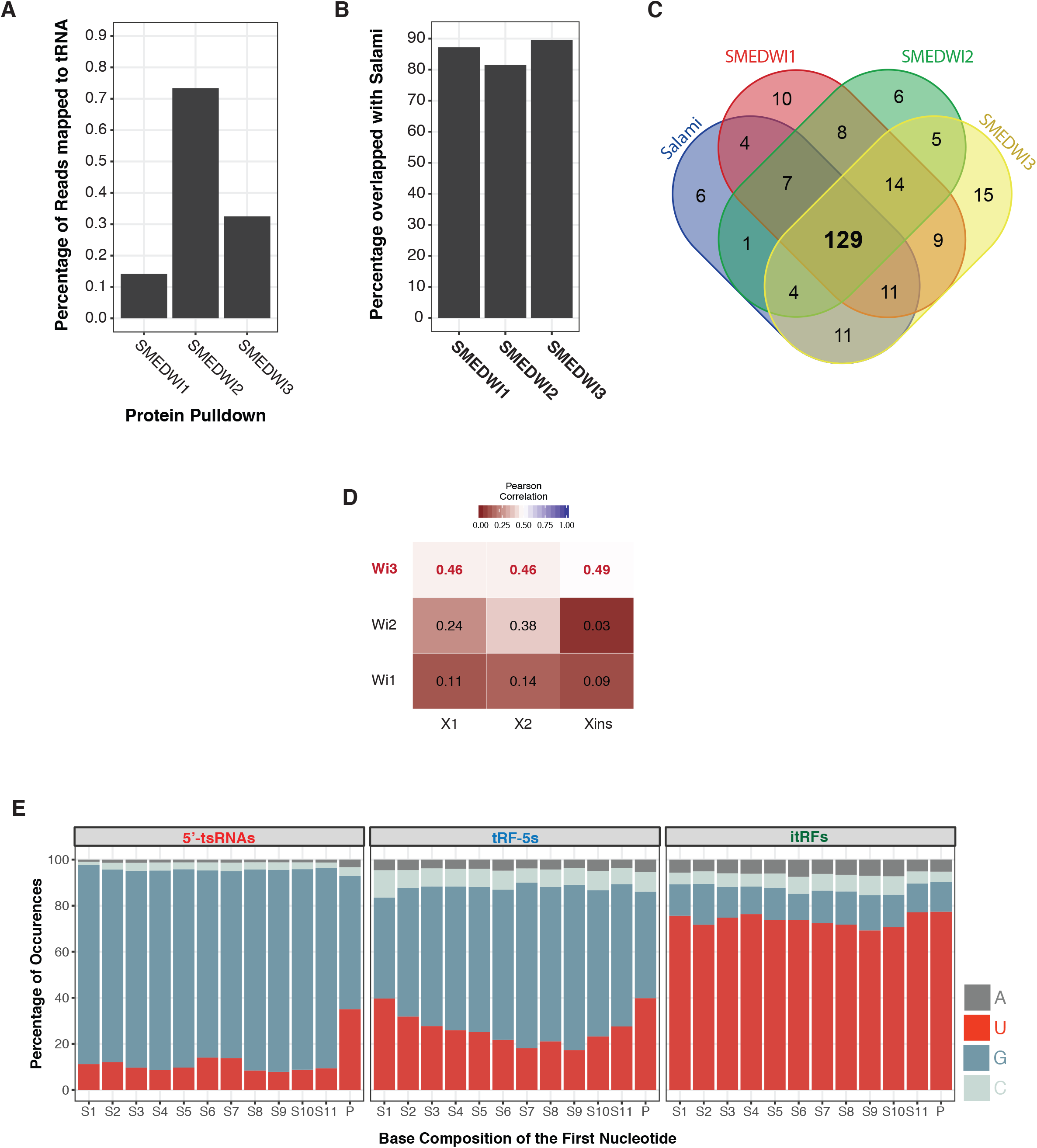
5’-tsRNA interaction with Piwi and sequence signatures in tRNA-derived fragments. A) Percentage of SMEDWI-1, −2 and −3 interacting RNAs that map to tRNAs. B) Percentage of identified planarian 5’tsRNAs associating with SMEDWI proteins in planaria. C) Pie chart of the common 5’-tsRNAs identified by salami sections and the three SMEDWI-associating 5’-tsRNAs. D) Correlation plot of 18-35 nt small RNA reads mapping to tRNA across SMEDWI-1, −2 and −3 with 18-35 small RNA reads mapping to tRNAs in X1, X2 and Xins cell population. E) Base preference of 5’tsRNAs, tRF-5s and itRFs at the first position, across planaria body sections. itRFs exhibit a strong enrichment for ‘U’.

### Sequence analysis of planarian tRFs indicate Dicer-based processing of itRFs

Understanding sequence features and base compositions have helped identify and characterize small RNAs across various systems (46). Sequence analysis of tRFs have provided valuable insights into the processing and functionality of tRFs. A ‘GG’ dinucleotide that is present across majority of tRNAs and thus retained in the tsRNAs have been shown to be important for the repressive role of tsRNAs in translation (47). Similar sequence analysis of 5’-tRFs identified TOG motifs (Terminal OligoGuanine) that facilitate G-quadruplex based interactions with Ybx1 (48). Interestingly, analysis of planarian tRF base composition revealed strong signature for ‘U’ at the first base of itRFs (Figure 7D and S7B). Similar preference for U among itRFs was observed across all the datasets used in this study (Figure S7C). This enrichment of ‘U’ at the first position was absent in tRF-5s and 5’-tsRNA that preferred a ‘G’ (Figure 7D and S7B). It is possible that this enrichment of ‘U’ could potentially arise because the parent tRNA might have an enrichment for ‘U’ at that particular position. To test this hypothesis, we first checked the start positions of itRFs on the tRNA and found that the majority of the reads originated from the 34^th^ position on the tRNA (Figure S7D). We next examined if all the tRNAs have an enrichment for ‘U’ at this (34^th^) position. Our analysis revealed that although there was a preference (~40%) for U at the 34^th^ position of the tRNA, we observed higher degree of enrichment (~70%) at the first position of all the itRFs (Fig 6D and S7E). Interestingly, positions 33 and 35 on tRNAs exhibited similar base preference for ‘U’, however very few itRFs were processed from these positions (Fig. S7D and S8E). This enrichment of U at the first position is characteristic of Dicer based cleavage (49). Dicer has already been implicated in the processing of tRFs in other systems and could also possibly be involved in processing itRFs in planarians (50). Further, the itRFs also display a more homogenous size distribution of ~20 nts that additionally suggests that itRFs may be processed by Dicer. Collectively our sequence analysis of the planarian tRFs reveals the possible involvement of Dicers in the processing of itRFs.

## Discussion

tRNA-derived fragments have been implicated in regulating various cellular processes through diverse mechanisms (2). However, most studies aimed at understanding tRFs have have been confined to a particular cellular process. Our study for the first time identifies and characterizes a role for these small RNAs in a complex biological process that is an embodiment of several cellular and molecular events, such as planarian regeneration. Due to lack of a comprehensive tRNA annotation in planarians, devising a stringent tRNA annotation pipeline was imperative to identify tRFs in planarians. Our pipeline identified 457 tRNA genes across *Schmidtea mediterranea* genome. These 457 tRNA genes can be broadly categorized into two groups (i) standard tRNAs that code for 20 standard amino acids (347) and (ii) Pseudo tRNAs (110). Interestingly, our prediction of tRNAs in planarians, failed to identify tRNAs for 7 anticodons. This is however a common occurrence as several genomes use near-cognate tRNAs to compensate in the absence of the correct codon; anticodon pair through wobble base pairing (51). Alternatively, the 7 unidentified anticodons could belong to the UNDET category of tRNAs. Surprisingly, one of the tRNAs exhibiting high gene copy number was a suppressor tRNA, SUP-TTA tRNA with 17 genes (spread across 52 genomic loci) in the planarians genome (Figure 1D). Suppressor tRNAs are mutants of standard tRNA genes that recognise a stop codon, thus delivers an amino acid to these positions instead of termination (25, 26). The large number of suppressor tRNAs in planarians, evokes an exciting possibility that these tRNAs could result in increased protein lengths thus altering their function. Understanding the function of these SUP-tRNAs role in translation regulation will add an additional layer of gene regulation in planarian biology. Our analysis also revealed a positive correlation between tRNA gene copy number and codon usage in planarians. Understanding the tRNA availability and codon has been important in shaping the translational landscape across organisms (52–54). The selection of the ‘optimal’ codons will be critical for optimizing heterologous expression of genes in the development of planaria transgenics.

Characterization of the tRNA-derived fragments in planarians, revealed three main classes of tRFs, the 18-24 nt species comprised of two species – the tRF-5s itRFs, and the dominant 25-35 nts 5’ tsRNAs. Expression analysis of these three species of tRFs across the planarian sections revealed 4 main clusters showing distinct spatial expression. A subset of 5’tsRNAs showed expression patterns similar to neoblasts-enriched transcripts suggesting that these 5’-tsRNAs could be key players in maintaining the stem cells homeostasis and differentiation. Further, our analysis also revealed certain 5’- tsRNAs, itRFs and tRF5s are enriched in the head and pharyngeal region while some are expressed throughout the planarian body. Our spatial expression analysis of tRFs adds to the catalogue of post-transcriptional regulatory mechanisms that exist in planarians. Analysis of tRFs during planarian regeneration revealed a divergent expression for all the three species suggesting that these three species could employ different modes of regulation during regeneration. It has been observed that 5’-tsRNAs largely influence the translation of transcripts, while the shorter 5’-tRFs and itRF may act akin to miRNAs, similar to what has been observed in *Drosophila* tRFs (4, 5, 55, 56). It would be interesting to understand the regulatory mechanisms these small RNAs employ to orchestrate planarian regeneration. Inspection of individual tsRNA expression revealed a dynamic change in expression during anterior regeneration (6 clusters) as compared to posterior regeneration (4 clusters). This observation could possibly be explained by the need to regenerate and reorganize the complex structures that make up the head as compared to the structures of the tail region. Our analysis also facilitated the dissection of ‘early’ and ‘late’ response tRFs. The ‘early’ tRFs could be essential for the wound closure and wound healing program whereas the ‘late’ tRFs could possibly regulate the differentiation and remodelling.

Our study also sheds light on the biogenesis and/or the modes of action of these small RNAs. Analysis of the parent tRNAs from which planarian tRFs are processed, revealed that a majority of these small RNAs are processed from specific sets of tRNA. Similar to higher organism, in planarians we observed Gly, Gln, and Asp were the most predominantly processed tRNA. It is interesting to note that this selective processing of tRNAs is evolutionarily conserved while our understanding of this process remains poor. 5’-tsRNAs (or tiRNAs or tRNA halves) have been shown to be produced by the endonucleolytic cleavage of tRNAs by angiogenin (3). Cleavage by angiogenin results in the tsRNAs carrying a 2’-3’ cyclic phosphates that are not amenable for conventional sequencing protocols (57). However, the data used in this study were obtained using conventional sequencing protocols. Moreover, we failed to identify a homolog for angiogenin in planarians. This strengthens our reasoning to believe that 5’-tsRNAs in planarians are processed in angiogenin-independent manner. However, studies in humans have suggested that tsRNAs could potentially be bound by PIWI (43). Our analysis of the recently published SMEDWI-1, −2 and –3 ClIP-seq data suggested that majority of the 5’ tsRNAs interact with PIWI proteins in planarians. It is interesting to note that SMEDWI-3, a piwi protein implicated to target coding transcripts, interacts with 5’ tsRNAs in all three cell compartments (X1, X2 and Xins). This evokes the possibility of PIWI targeting some of the coding transcripts identified in a previous study, mediated by the 5’-tsRNAs (44). Lastly, we observed a high base preference for ‘U’ among the itRFs suggestive of Dicer-based cleavage. It is known that planarians possess two Dicers homologs and is important for planarian regeneration (58, 59). However, it remains unclear if both these Dicers process miRNA and itRfs or they are mutually exclusive. It would be interesting to explore the effects of Dicer and Ago knockdown on the production and function of the itRFs.

In conclusion, we present the first-ever report and characterisation of planarian tRFs using high-quality tRNA annotations. The regulatory roles of tRFs remain poorly understood at an organismal level and our characterization of these small RNAs in planarians solidifies the notion that tRFs are an important family of small RNAs with impactful regulatory roles across all life forms. Further, the dynamic expression of planarian tRFs during regeneration suggests active roles for these small RNAs in complicated biological processes. Finally, our study brings to the fore, a previously unknown layer of regulatory complexity to the process of planarian regeneration and promises to be a new avenue of research in the quest to understand the process regeneration. Moreover, molecular tractability of planarians makes them an ideal invertebrate model system to explore the function and biogenesis of these small RNAs *in vivo*.

## Materials and Methods

### Predicting putative tRNA gene families

We used dd_Smes_G4 (21) assembly of *Schmidtea mediterranea* genome to identify putative tRNA genes. We used two programs to identify tRNA gene loci across planarian genome: tRNA-SCAN-SE (version 1.3.1) and Aragorn (60, 61). We enabled intron prediction for both the programs and used the following parameters to predict: tRNA-SCAN_SE (*-G -y -l -o -f -m -p*) and Aragorn (*-t -gcstd -I -seq -br -fasta -o*). Both the programs predicted ~4100 sites across planarian genome (4115 – tRNA-SCAN-SE and 4143 – Aragorn). We clustered the sequences predicted by both the programs using CD-HIT with 90% sequence similarity cutoff and identified 708 unique sequences (62). Among 708 sequences, 302 were predicted by both the programs, remaining 406 sequences (132 and 274) were predicted only by tRNAScan-SE and Aragorn respectively. We further removed false positives from these 708 predicted sequences using tRNA-SCAN-SE (version 2.0.5, *-o -f -m -a -l -p -detail -y –isospecific –thread*) (23). This improved version of the algorithm is known to have incorporated methodologies with improved probabilistic search and gene models. We finally narrowed down 457 unique sequences which could code putatively for tRNA genes. We used VARNAv3-93 for obtaining secondary structures of tRNA (63) as shown in Figure 1E.

We further used scores derived from TFAM 1.0 classifier (*-E -t -s*) to assign an amino acid to the UNDET tRNAs identified from our method (24). Using TFAM classifier we identified three initiator tRNA (*iMet*) in planaria genome (Figure 1 and Table S1). iMet tRNA was identified based on sequence features highlighted in Figure 1E and isotype specific score from tRNA-SCAN-SE(v2.0.5). We also aligned predicted planarian tRNA sequences codon wise using MacVector and given as extended supplementary figure1.

### Codon usage calculation

To calculate codon usage in planaria, we downloaded the latest version of transcriptome annotation (SMESG – repeat filtered) that has 30,917 genes (~59,800 isoforms) from planmine (36). We only considered ORFs which are of length >= 100 and <= 10,000 nucleotides. We also removed the gene sequence which has Ns in the ORF regions. Codon usage is the measure of frequency of occurrence of all the possible three letter codon in coding transcripts (ORFs). We wrote a custom made perl script to calculate this parameter.

#### Frequency per 1000 codons

This index, is a ratio of occurrence of each codon to the total number of triplets from all coding ORFs and normalized per 1000 codons. Frequency per 1000 codons gives the global profile of codon usage for a particular organism (Table S2).

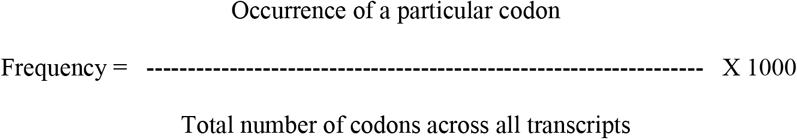

Percentage of tRNA loci per codon is calculated similar to codon fraction for each codon.

### Small RNA datasets in planaria

We downloaded the publicly available small RNA datasets of planaria from NCBI-SRA. Planarian cell population (X1,X2 and Xins) and regeneration time point data was downloaded from SRA065477 (64). Smed-piwi pulldown data was downloaded from GSE122199 (44).

### Identification of tRNA-derived fragments in planaria

We used dd_Smes_G4 (21) assembly of *Schmidtea mediterranea* genome and tRNAs predicted from this study for analysis. Small RNA sequencing data downloaded from SRA were converted to fastq files using sra-toolkit (https://ncbi.github.io/sra-tools/). From the sequencing reads, we trimmed TruSeq small RNA adapters using cutadapt program (-f fasta -b TGGAATTCTCGGGTGCCAAGG -O 5 -m 6 -o) (65). Adapter trimmed reads were aligned to flatworm (planarian) rRNA sequences retrieved from NCBI and unmapped reads were used for further analysis. We downloaded planarian miRNA sequences from miRbase (66) and piRNA sequence coordinates from Kim *et al.*, paper. For analysis, we considered reads ranging from 18 and 35 nucleotides and mapped these to genome and other databases using bowtie v1.1.2 (*-f -v 2 -p 20 –un*)(67). We used two mismatch as a constant parameter for mapping all the reads used in this study, as we did not observe much deviation in our results with varying mismatches ranging from 0 to 2. To calculate the percentage tsRNA reads that mapped to genome, we calculated the ratio of reads of a particular size that mapped to tRNA to the total reads that mapped to the genome. Further, we calculated per base tRNA coverage using the following formula given below. We used the coverage values obtained to plot the tRNA coverage heatmaps and area plots. We used customized perl script for all the analysis used in this study. We used R ggplot2 library for plotting (68). We followed a similar analysis pipeline as described in (7).

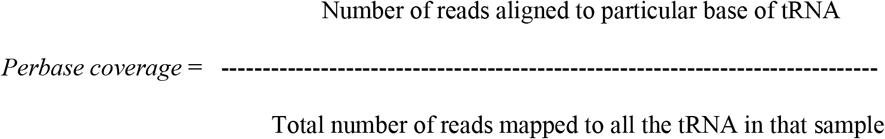

### Small RNA counts and data normalization

We obtained raw read counts mapping to individual tRNA using customized perl script. We used the well-established DESeq algorithm for normalization of the sequencing data and identification of differentially expressed tRFs (adj. *P*-value < 0.05) (69). All the statistical tests were done in R (70). The normalized values obtained from DESeq is used for clustering the tsRNAs based on expression across multiple datasets. Top ten candidates were represented as percentages and plotted in a pie-chart as shown in Figure 2D. We used MFuzz (R package), and soft clustered expression values across multiple timepoints to obtain dominant cluster patterns (71). All the heatmaps depicting expression changes are plotted using a R package pheatmap, (https://cran.r-project.org/web/packages/pheatmap/index.html).

### Small RNA sequencing from salami sections

Asexual planarians were killed and sliced into 12 sections as described in (37). The sliced pieces were put in Trizol to isolate RNA. Small RNA libraries were made using Illumina TruSeq Small RNA Prep kit and later sequenced on Nextseq500 machine. We analyzed the data as described above in the methods. We averaged the normalized value derived from each fragment to get an estimate of expression in whole animal.

### *Smedwi* pulldown data correlation

We downloaded smed-PIWI protein pulldown data from GSE122199 and processed it to identify putative tRFs as described in the methods above. We normalized the individual tRFs using DESeq. To draw correlations between piwi-associating tRFs and planarian cell populations, we used small RNA sequencing data from X1,X2 and Xins population from SRA065477. Small RNA sequencing data was processed as described above in methods and normalized value from three distinct cell population is correlated with the pulldown data (Table S10). We calculated pearson correlation and plotted as matrix using R and ggplot2 module.

### Base preference in small RNA reads

We categorized reads mapping to the three identified pools (tRF5s, itRFs, tsRNAs) based on read length and position in which reads align to an tRNA. We then size segregated reads which fall under these categories and calculated per-base preference using custom perl script. Perbase coverage is calculated as occurrence of A,T,G,C at each position divided by total number of reads under each category. The derived percentages were later plotted using R ggplot2 package.

### Understanding dynamic processing of tRNAs

From the sequencing data, we observed three distinct pools of tRNA-derived small RNA fragments in planaria. To check if there are any overlap in tRNAs that are processed in generating these fragments (tRF-5s, itRFs and tsRNAs) we devised this following strategy. To assign if a particular tRNA is giving rise to one specific pool, it has to satisfy these three criteria: (i) >90% of reads mapping to that particular tRNA should map either before (5’ end) or after (3’ end) anticodon position. This is relative to the fragments that is assessed. For example, if the fragment is itRFs, >90% of reads mapping to that particular tRNA should map after the anti-codon (3’end). (ii) Remaining 10% reads, should be less than 10 reads as it would be negligible amount to define it as one of the pools, and (iii) Since the three identified species is of varied lengths (tRF5s, itRFs – 18-24nt and tsRNAs – 25-35nt), we made sure that the expression of third species is less than median normalized value and has fourfold lesser reads mapping. For example, if the fragment is itRFs (18-24nt of length), at least fourfold lesser number of reads (w.r.t 18-24nt reads mapping at 3’end) of length 25-35nt should map to 5’ end of tRNA. The number of 25-35nt reads mapping at 5’end should be less than median normalized value. Based on these criteria’s we classified tRNAs into capable of coding all three or either of two or specific to one pool of tRNA derived small RNA fragments. Number of tRNAs that are categorized is shown in Fig.S3 E.

### Northern hybridizations

The RNA blot was performed as described previously (72, 73). 10 μg of total RNA was isolated from whole planaria (homeostasis) and resuspended in 8 μl loading buffer (0.10% bromophenol blue, 0.10% xylene cyanol in 100% de-ionized DEPC-treated formamide), heated at 95°C for 1 min, and loaded on to a 15% denaturing polyacrylamide gel (a 19:1 ratio of acrylamide to bisacrylamide, 8 M urea). The gel was run at 100 V for 3 h and then transferred to a Hybond N+ membrane by electroblotting at 10 V overnight at 4°C. After UV crosslinking (UVP), hybridization was performed at 35°C for 12 h in UltraHyb-Oligo buffer (Ambion) containing desired probes (*Pseudo_GlyGCC_10* – TACCACTGAACCACCAATGC; *UNDET-Gly_17* – TACCACTGAACCACCGATGC; *UNDET-Gln_23* – ACGCCTACACCATGGACCTC; *GlyTCC_6* – GACCGTTACACCACAATCGC; *Asn-GTT_5* – AATTGCGCCACGGAGGCTC; *iMet-CAT_1* – TCCACTGCGCCACTCTGCT; *SUP-TTA_2* – CCGCTTACACCATCGAACC). DNA oligos complementary to candidate 5′-tsRNAs, tRF-5s, itRFs were end-labeled with ^32^P-ATP (Board of Radiation and Isotope Technology, India) using polynucleotide kinase (NEB), purified through MicroSpin G-25 Columns (GE Healthcare), and were used as probes. The blot was washed twice with 2× SSC, 0.5% SDS for 30 min at 35°C. The signal was detected after exposure on a phosphorimager screen using a Molecular Imager (GE Healthcare). All the tRF candidates were analysed in five biological replicates.

### Planaria culture

Animals used in this study belong to the sexual strain of species *Schmidtea mediterranea*. They were maintained at 20°C in planarian media (2 mM NaCl, 0.1 mM KCl, 0.1 mM MgSO_4_, 0.12 mM NaHCO_3_ in distilled water) and fed beef liver paste. Animals were starved two week prior to experiments.

## Data Availability

The small RNA sequencing data from planarian salami sections generated as a part of this study is deposited in NCBI-SRA under the project id SRP277000 (PRJNA646861).

## Author Contributions

SK supervised the study. SK, DP and VL, conceived and designed the study. VL performed all the analysis with extensive inputs from SK. DB performed the salami sectioning of planaria. STN and PVS contributed to Northern Hybridizations. SK and VL wrote the manuscript with inputs from DP. All authors have read and approved this manuscript.

## Acknowledgements

We thank the members of Palakodeti and Gulyani lab at inStem for critical comments and discussions. We extend our gratitude to Jochen Rink for hosting and training DB on salami sectioning. We thank the inStem-NCBS Next Generation Genomics Facility for catering to the sequencing needs. We acknowledge the authors of the previous studies whose data has been used in this study. We extend their thanks to Nishan Shettigar for critical comments on the the manuscript. VL is supported by CSIR-SRF and this work was funded by DST Swarnajayanti Fellowship (DST/SJF/LSA-02/2015-16) awarded to DP.

## Declaration

The authors declare no conflict of interest.

## Notes

### Competing Interest Statement

The authors have declared no competing interest.

