## Supplementary Figures S1 - S7 for "Comprehensive annotation and characterization of planarian tRNA and tRNA-derived fragments (tRFs)"

#### Supplementary Figure S1

**A**

| | tRNA genes | Genomic Loci |
|---|---|---|
| 20 Standard amino acids | 303 | 942 |
| UNDET tRNAs | 44 | 125 |
| Pseudo tRNAs | 110 | 305 |

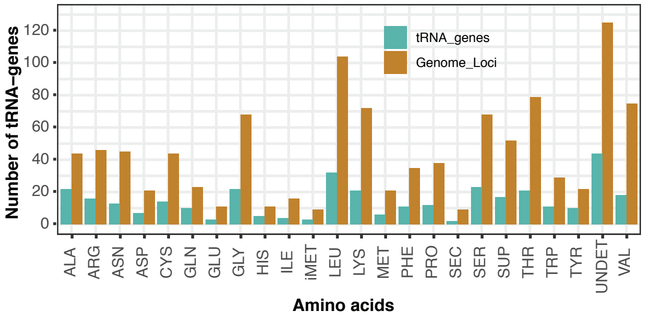


**B**

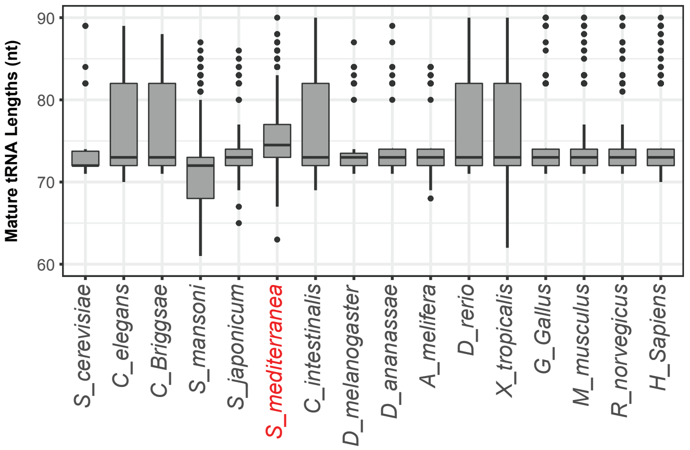


**C**

**tRNA (anticodon) Number of Genes**

**Selenocysteine-tRNA Sec- TCA** 2 **Undetermined-tRNA (Undet)** 44

| First Base | Second Base: A | Second Base: G | Second Base: U | Second Base: C | Third Base |
|---|---|---|---|---|---|
| A | AAA 6 10 (Phe) | AGA 7 2 | AUA 2 1 (Tyr) | ACA 3 0 (Cys) | A |
| G | GAA 5 2 | GGA 0 1 (Ser) | GUA 8 1 | GCA 11 3 | |
| U | UAA 7 4 (Leu) | UGA 7 2 | UUA 11 11 Sup | UCA 3 0 Sup | |
| C | CAA 4 1 | CGA 4 4 | CUA 3 1 | CCA 11 3 (Trp) | |
| A | AAG 4 2 | AGG 3 2 | AUG 0 0 (His) | ACG 2 1 | G |
| G | GAG 0 0 (Leu) | GGG 1 0 (Pro) | GUG 5 2 | GCG 1 0 (Arg) | |
| U | UAG 11 2 | UGG 7 1 | UUG 8 0 (Gln) | UCG 2 0 | |
| C | CAG 6 1 | CGG 1 1 | CUG 2 1 | CCG 1 1 | |
| A | AAU 2 2 | AGU 5 1 | AUU 2 0 (Asn) | ACU 3 0 (Ser) | U |
| G | GAU 0 0 (Ile) | GGU 2 0 (Thr) | GUU 11 1 | GCU 2 0 | |
| U | UAU 2 0 | UGU 10 2 | UUU 15 9 (Lys) | UCU 6 0 (Arg) | |
| C | CAU 6 2 (Met) / 3 0 (iMet) | CGT 4 0 | CUU 6 2 | CCU 4 1 | |
| A | AAC 6 0 | AGC 6 0 | AUC 0 0 (Asp) | ACC 0 0 | C |
| G | GAC 0 0 (Val) | GGC 4 3 (Ala) | GUC 7 3 | GCC 10 4 (Gly) | |
| U | UAC 4 1 | UGC 7 2 | UUC 1 1 (Glu) | UCC 6 0 | |
| C | CAC 8 0 | CGC 5 0 | CUC 2 1 | CCC 6 0 | |

**Figure S1: A)** Table showing total number of tRNA genes predicted in planaria and the number of genomic loci for these tRNAs. **B)** Mature tRNA lengths across different species shows a median length of 72 nucleotides with planarian tRNAs showing a median of 75 nts. **C)** tRNA anticodon table decpicting all the anticodons and the number of tRNA genes identified for each anticodon (in green) and pseudogene (in grey).

#### Supplementary Figure S2

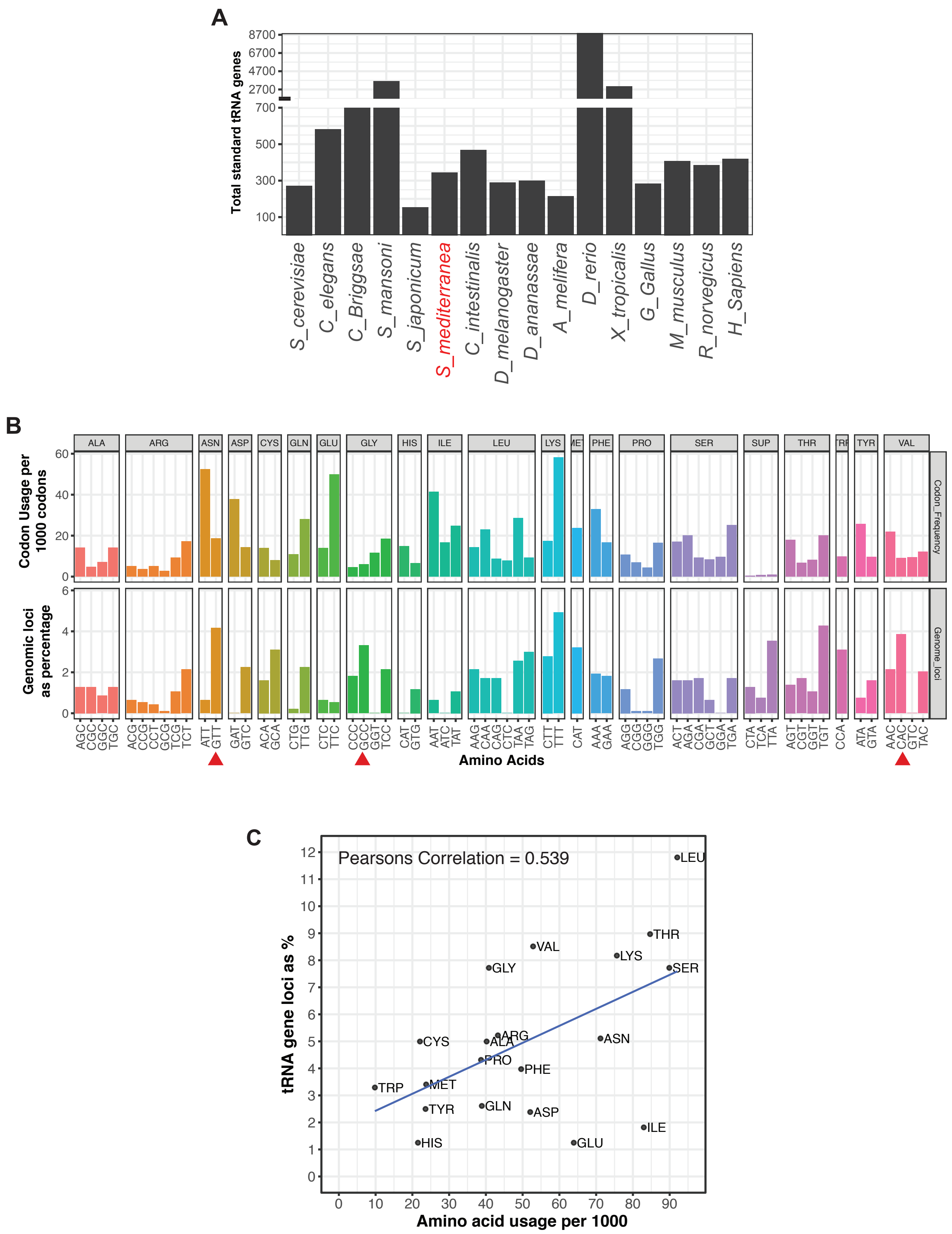


**Figure S2: A)** Number of tRNA genese for the standard 20 amino acids across the animal kingdom. Predicted planarian tRNA gene numbers are comparable across the species. **B)** Codon frequency across 1000 codons in the transcriptome and the abundace of its cognate anticodon tRNA in S.mediterranea genome. The red arrows examples of codon:anticon pairs that show opposite correlation. **C)** Correlation between tRNA gene copy number and amino acid usage in planarians.

### Supplementary Figure S3

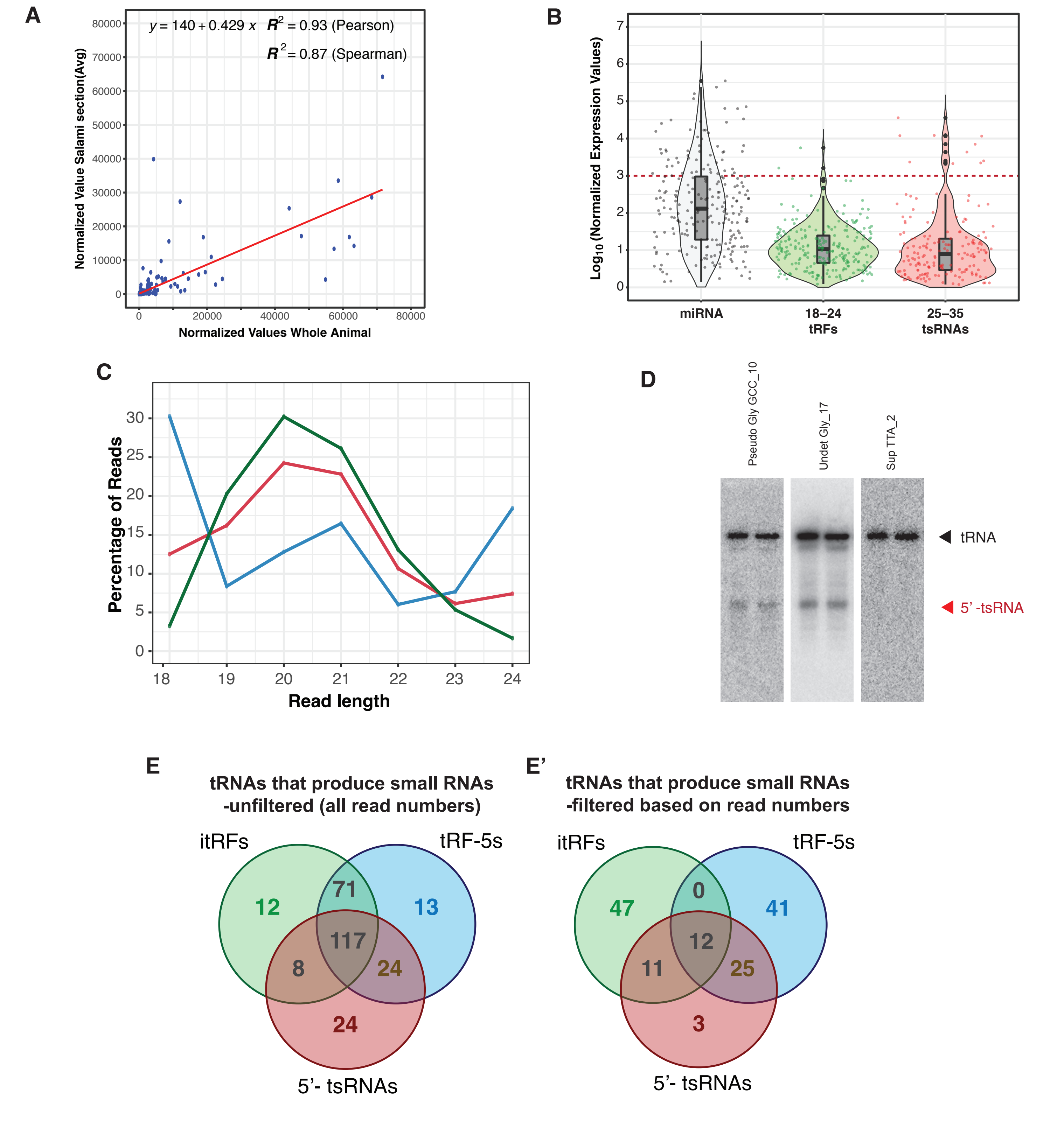


**Figure S3: A)** Correlation between small RNA reads mapping to tRNA in 'Uncut' and the cumulative of small RNA reads mapping to tRNA from Salami Sections. **B)** Normalised expression of all miRNAs compared to the reads mapping to tRNAs. The red dotted like indicates that the top 20 25-35 nt tsRNA species are expressed at higher levels than most miRNAs. **C)** Size distribution of itRFs and tRF-5s and the over all 18-24 reads mapping to tRNA. **D)** Northern blot against top enriched 5' -tsRNAs (Pseudo-GlyGCC and UNDET-Gly) in planaria shown in replicates. Sup-TTA is used as a control which does not generate 5'-tsRNAs in planaria. The black arrow marks the parent tRNA and the red arrow marks the 5'-tsRNA. **E)** Venn diagram of tRNA that prodice itRFs, tRF-5s and 5'-tsRNAs. **E')** Venn diagram of tRNA that prodice itRFs, tRF-5s and 5'-tsRNAs using stringent criteria.

### Supplementary Figure S4

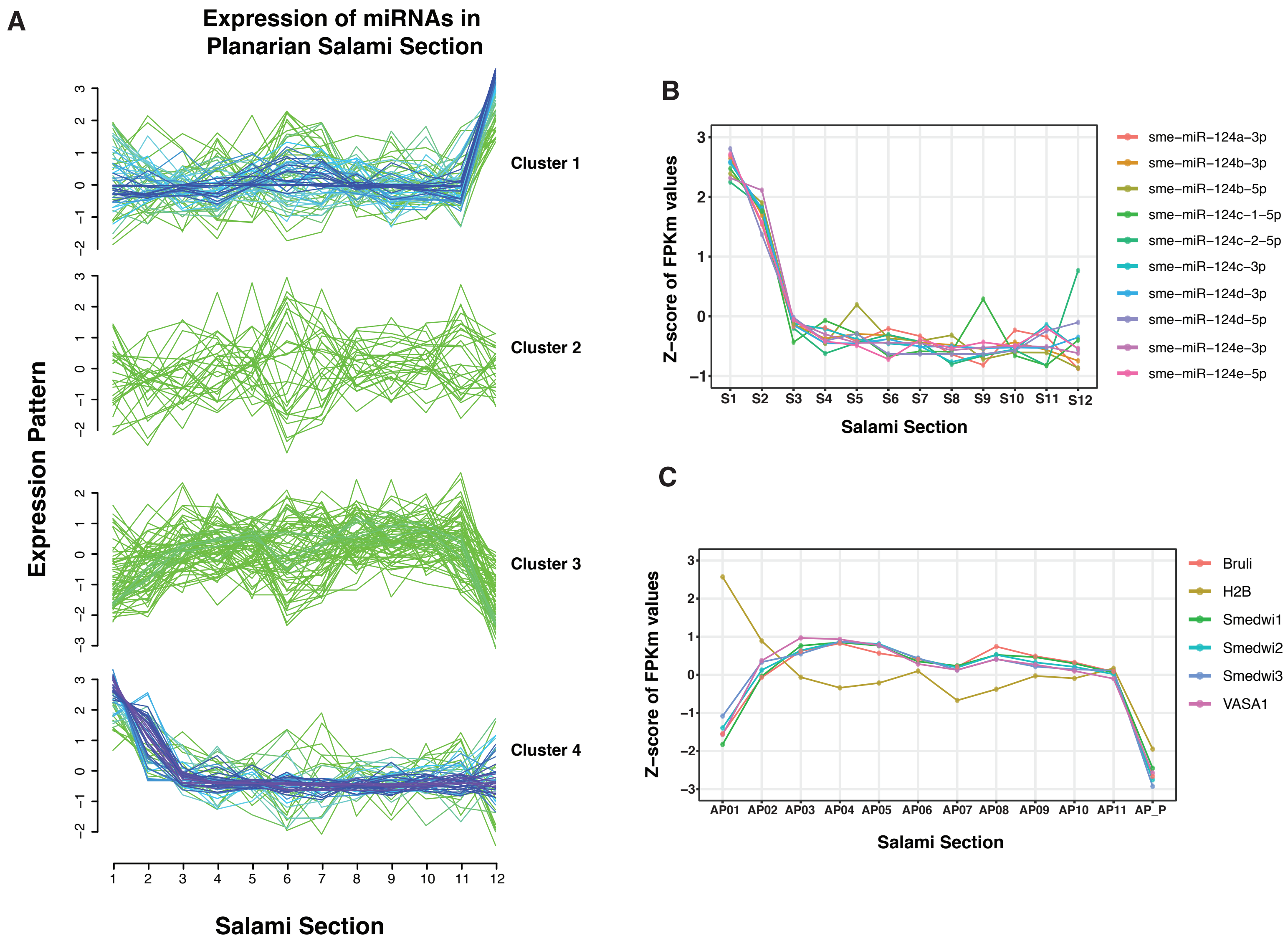


**Figure S4: A)** Expression clusters of miRNAs across planarian body axis identifies 4 clusters of expression. **B)** Expression of miR-124 family of miRNAs that are expressed in the brain shoes enrichments in the anterior sections of planaria. **C**) Expression of transcripts expressed in neoblasts across the different salami sections.

### Supplementary Figure S5

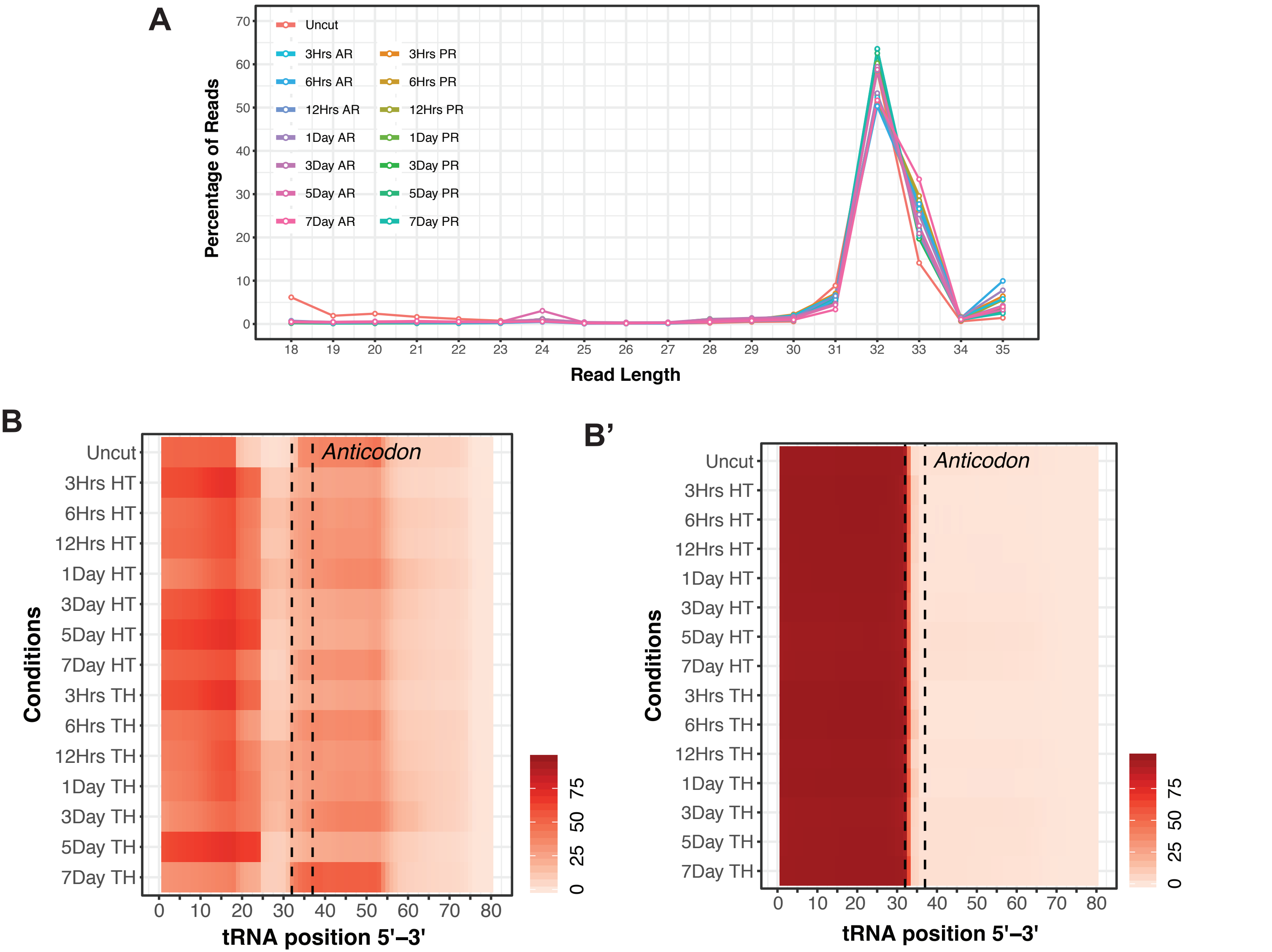


**Figure S5: A**) Size distribution of 18-35 nt small RNA reads that map to planarian tRNAs across regeneration timepoints. **B)** Perbase coverage of 18-24 nts of small RNA reads mapped to the tRNA length identifies two species of tRNA fragments - itRFs and tRF-5s. **B')** Perbase coverage of 25-35 nts of small RNA reads mapped to the tRNA length shows reads arise only from the 5' arm of tRNA.

### Supplementary Figure S6

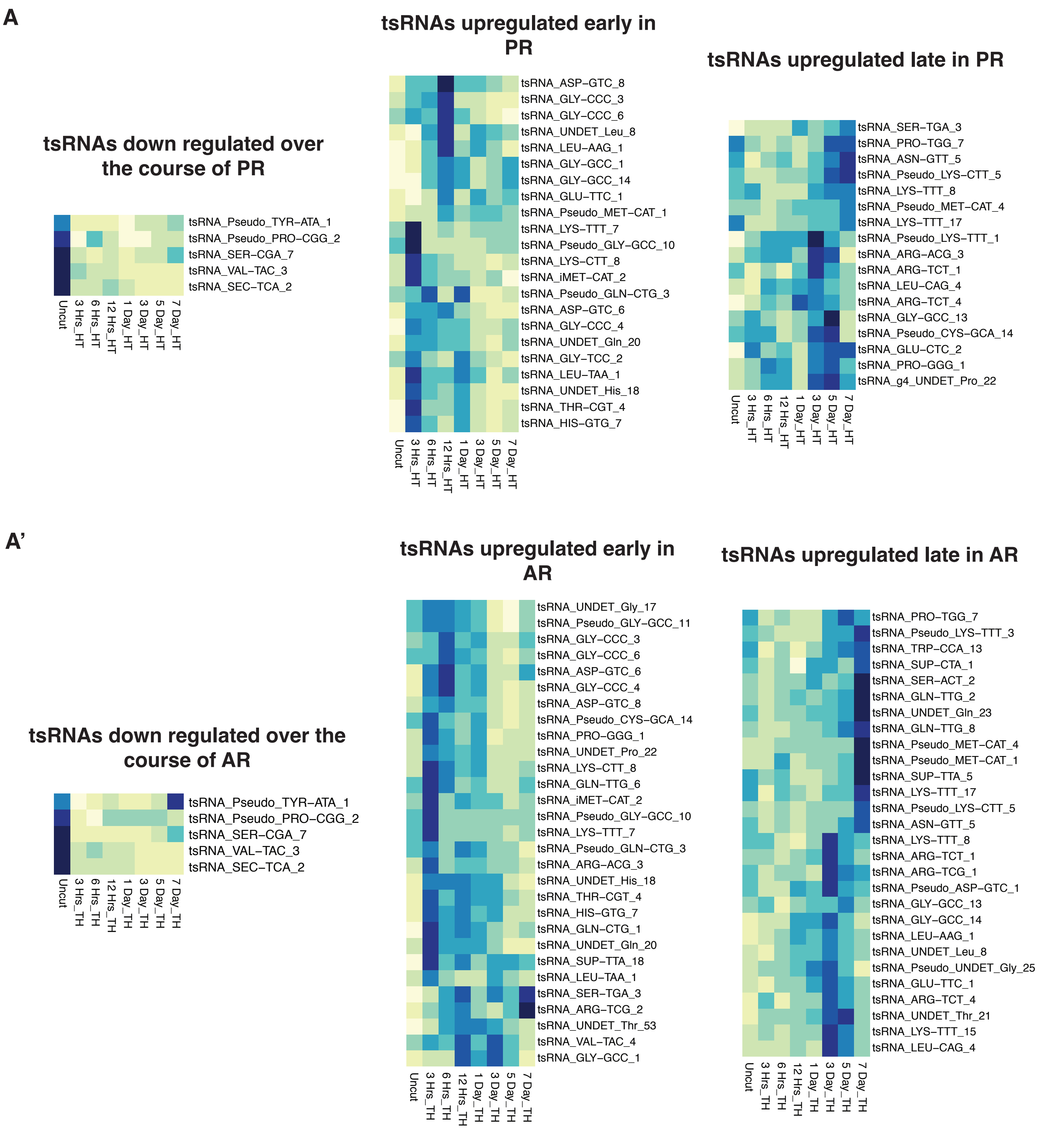


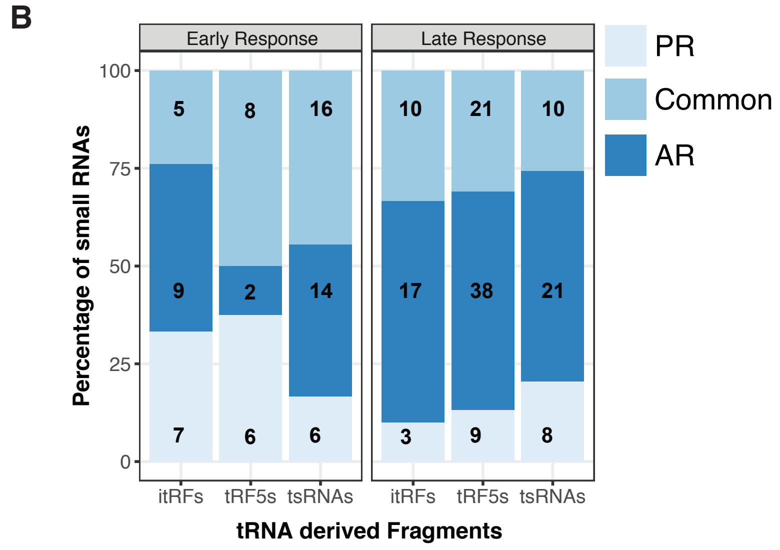


**Figure S6: A**) Heatmaps of 5' tsRNAs during posterior regenerating (PR). **A')** Heatmaps of 5' tsRNAs in anterior regenerating worms (AR). **B)** Percentage of tRNA-derived fragments that are common and unique in early (3hrs -1day) and late (3days - 7days) regeneration of head (HT)and tail fragments (TH).

### Supplementary Figure S7

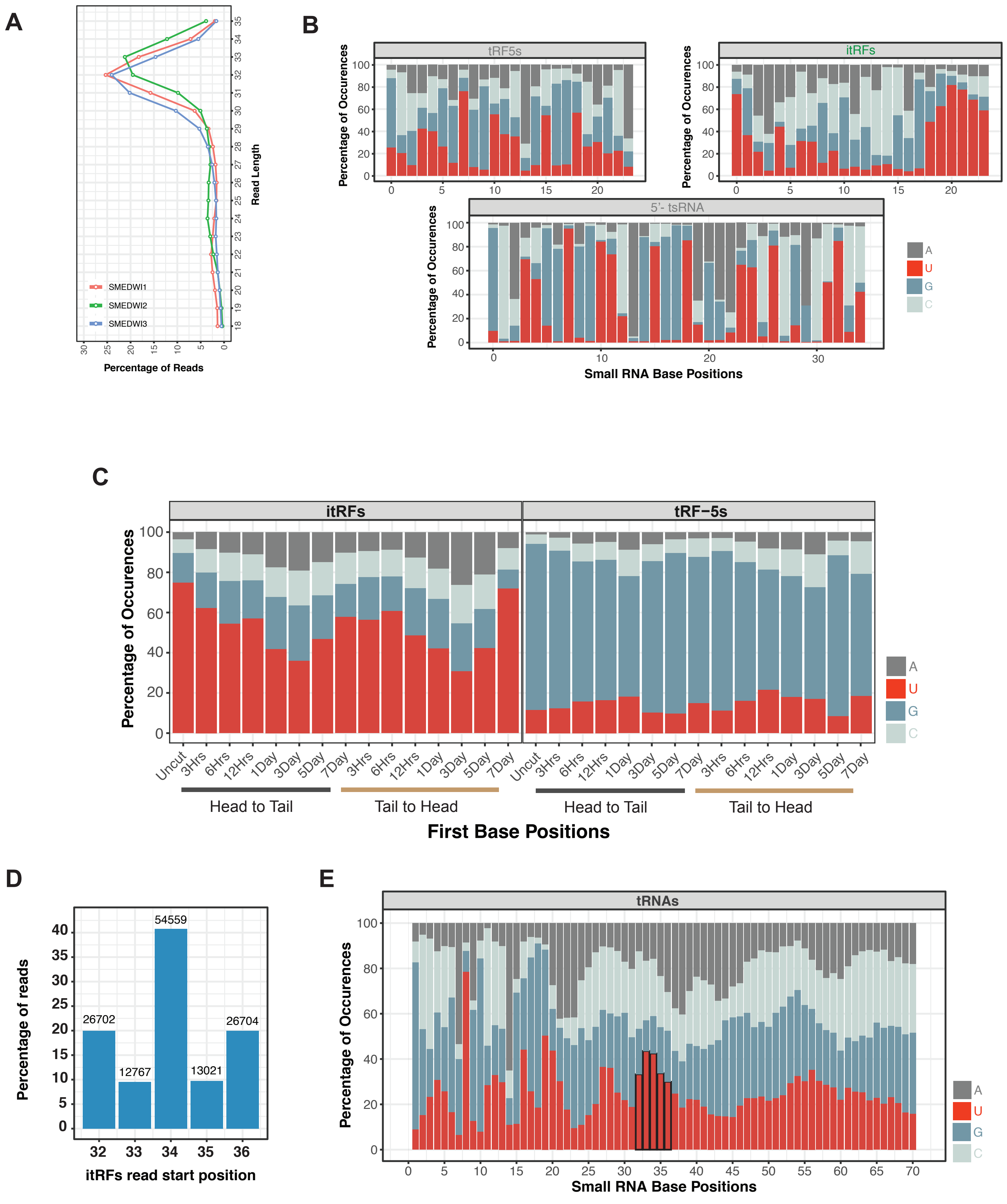


**Figure S7: A**) Size distribution of reads associating with SMEDWI-1, -2 and -3 that map to tRNAs. **B)** Base preference for the first base of all itRfs and tRF-5s during regeneration timepoints. **C)** Base preference across the length of tRNA-derived fragments identified in the study. **D)** Start positions of itRF fragments on the respective tRNAs from which they are processed. **E)** Base preference for the positions (from which itRFs are processed) across all tRNAs.
