## Extended Supplementary 1 for "Comprehensive annotation and characterization of planarian tRNA and tRNA-derived fragments (tRFs)"

### Formatted Alignments

```
                          10         20         30         40         50         60         70         80         90         100        110        120
dd_Smed_g4_ALA-AGC_tRNA_1 --------------GGGGAACGTAGCTCACATGGTAGAG-C-GCTCGCTTAGCATGCGGAAGAGGTAC---TGGGATCGATACC-C-AGGTTCTCCA-----
dd_Smed_g4_ALA-AGC_tRNA_2 --------------GGGGGTATAG-CTCAGTG-GTAGAGGCATGTCGACTAGCAGA-TCGCAGAAGTC---CCGGTTCAAATCC--GGGTGCCCCCT-----
dd_Smed_g4_ALA-AGC_tRNA_3 --------------GGGGAACTTAGCTCACATGGTAGAG-C-GCTCGCTTAGCAAT-GCGAGAG-TAC---TGGGATCGATACC-CCAGGTTCTCCA-----
dd_Smed_g4_ALA-AGC_tRNA_4 --------------AGGGAACT-AGCTCACATGGTAGAG-C-GCTCGCTTAGCAATAGCGAGAGGTA----TGGGATCGATACC-CAGGTTCCTCT------
dd_Smed_g4_ALA-AGC_tRNA_5 --------------GTCGTGAATGGCCGAAGTGGTTAAG-GCGTTGGACTAGCAAATTCCAAATGGGTT-CCCCGCGCAGGTTCAAATGCCTGCTCACGACG
dd_Smed_g4_ALA-AGC_tRNA_6 --------------GGGGAACT-AGCTCACGATGGAGAG-C-GCTCGCTTAGCA-T-GCGAGAGGTAC---TGGGGATCGTACC-CCAGGTTCTCCA-----
dd_Smed_g4_ALA-CGC_tRNA_1 -----------------GAGCTCGTGGCGCAACGTTAGC-GCGTCTGACTCGCAGATCAGAAGAAGGTT-GCGTGTTCGATCACGTCGGGCTCA--------
dd_Smed_g4_ALA-CGC_tRNA_2 --------------GGGGATGTAAGCTTCAGTGGTAGAG-CGTTCCGCTTCGCAAT-GTGGAAAGGTC--CCGGGTTCGATCCC-C--GCATCTCCA-----
dd_Smed_g4_ALA-CGC_tRNA_3 -----------------GAGCTCGTGGCGCAACGTTAGC-GCGTCTGACTCGCAGATCAGAAGAAGGTT-GCGTGTTCGATCACGTCGGGCTCA--------
dd_Smed_g4_ALA-CGC_tRNA_4 --------------GCCCCTATAG-CTCAGAAGGTAGAG--CACTGTCCTCGCT---AAACAGGGGTCG-AGACGTTCACTTCTCTCTGGGGGCA-------
dd_Smed_g4_ALA-CGC_tRNA_5 --------------GGGGATATAG--CTCAGTGGTAGAG-C-GTTCGCTTCGCA-T-GTGAAAGGTCC---CGGGTTCAATCC--CCGGTATCTCCA-----
dd_Smed_g4_ALA-GGC_tRNA_2 --------------TCCTCGATAGTTAAAGTGGGTCAGTAATCTCCGCCTGGCA--CGTGGAAAGCCC----GGGTTCGATTCC--CGGTCGGGGAG-----
dd_Smed_g4_ALA-GGC_tRNA_3 --------------GGGGGTATAG-CTCAGTG-GTAGAG-CATTCCGACTGGCAGA-TCGAGAAGGTC---CCCGGTTAAATCC--GGGTGCCCCCT-----
dd_Smed_g4_ALA-GGC_tRNA_6 --------------GTCGTGA-TGGCCGA-GTGGTTAAG-GCGTTGGTTAGGCGAATCCAATGGGGTTTTCCCCGCGCAGGTTCAAAT-CCTGCTCACGACG
dd_Smed_g4_ALA-GGC_tRNA_7 --------------GGGGGTATAG-CTCAGTG-GTAGAG-CATTCCGACTGGCAGA-TCGAGAAGGTC---CCCGGTTAAATCC--GGGTGCCCCCT-----
dd_Smed_g4_ALA-TGC_tRNA_1 GGGGATGGTAGCTCAGTGGTAGAGCTTCGTCTGGTAGGC-GGTTTCGCTTTGCA-T-GTGAAA-GGTC--CCGGGTTCGAATCC-CCGGCATCTCCA-----
dd_Smed_g4_ALA-TGC_tRNA_2 --------------CGCTCGATAG-CGCAGTAGGTAGCG-CAGTCAGTTTTGCGTCATTGAAG---TCG--TGAGTTCGATTCC-TCACTCGGGCAT-----
dd_Smed_g4_ALA-TGC_tRNA_3 --------------GGGGATGTAGCACTCAGTGGTAGAG-CCGTTCGATTTGCA-T-GTGAAAAGGTC--CCGGGTTCGATCCGGCCGGCATCCTCA-----
dd_Smed_g4_ALA-TGC_tRNA_4 --------------GGGGGTATAG-CTCAGTGCATAGAG-CATTC-GACTTGCAGA-TCGAGAGGTCC---CCGTTTCAAATCC--GGGTGCCCCCT-----
dd_Smed_g4_ALA-TGC_tRNA_5 --------------GGGGATGTAG-GCTCAGTGGGAGAG-C-GTTCGCTTTGCAAT-GTGAAA-GGTCGTCCCGGTTCGATCC--CCGGCATCTCCA-----
dd_Smed_g4_ALA-TGC_tRNA_7 --------------GGGGATGTAGCTGCAAGTGGTAGAG-C-TTTCGCTTTGCA-T-GTGAAAGGTCC--CCGGGTTTCGATCC-C--GCATCTCCA-----
dd_Smed_g4_ALA-TGC_tRNA_9 ----------------GGATGTAGCTTCAAGTGGTAGGC-GGTTTCGCTTTGCA-T-GTGAAAGGT-C--CCGGGTTCGAATCC-CCGGCATCTCCA-----
                          GGGGATGGTAGCTCGGGGATMTAGSCTCARDTGGTAGAGRCMGTTCGCTTWGCARWHGYGAAAGGGHCTTCCGGGTTCGATTCCRCCGGYMTCYCCACGACG
```

### Formatted Alignments

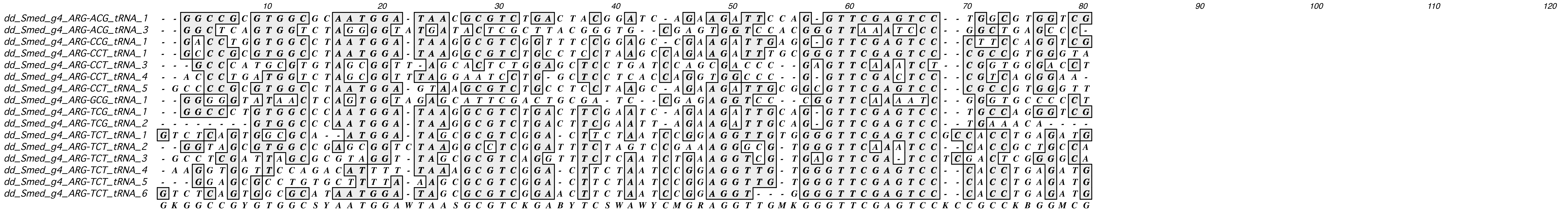

### Formatted Alignments

```
                                  10        20        30        40        50        60        70        80        90       100       110       120
dd_Smed_g4_ASN-ATT_tRNA_1      -GCCTCGACTAGCGCAGTAGGGTAG-GCGCGTCGTCTATTATATGAA--TGGTCGTGAGTTCG---ATCCTCA-TCGGGGCA----
dd_Smed_g4_ASN-ATT_tRNA_2      ----GCTAGGATGGCCGATGGTTAAGGCGGTGGACTTATTAAGTCCA-CTGGACAATGTCCCGTGGGTTCGAACCCACTTCTAGCA
dd_Smed_g4_ASN-GTT_tRNA_1      ---GGCGTGATGTCCTAGTGGTTAGGGA-ATTGCGTTGTTGGCCGCA--ATAACCCAGTTTCG---AATC-TGGTCACGCCA----
dd_Smed_g4_ASN-GTT_tRNA_10     ---GGCGTGATGTCCTAGTGGTTAGGGA-ATTGCGTTGTTGGCCGCA--ATAACCCAGTTTCG---AATC-TGGTCACGCCA----
dd_Smed_g4_ASN-GTT_tRNA_12     -ACGGTG-A-AGCTCAGTTGGTAGAGGGATCAGGACTGTTATTTCCTGTAGGTCGGTGGTTCA--AATCCGCC-TCACCGGA----
dd_Smed_g4_ASN-GTT_tRNA_13     TTTATTG-ATAGCTCAGTTGGT--AGAGCTGAGGACTGTTGATCCTT--AGGTCGGTGGTTCA--AATCCGCC-TCACCGGAA---
dd_Smed_g4_ASN-GTT_tRNA_2      -GCCTGCCGTGGCGCAATTGGTTAGCGCGTT-CGCTTGTTTCCGCAA--AGGTTGGTGGTTCG---AGCC-AC-CCGGGGCG---
dd_Smed_g4_ASN-GTT_tRNA_3      -GCCT-CCGTG-CGCAATTGGTTAGCGCGTTACGGCTGTTAAACGAA--AGGTTTGTGGTTCG---AGCCCAC-CCGGGGCG---
dd_Smed_g4_ASN-GTT_tRNA_4      -CCGGTG-ATAGCTCAGTTGGTAGAGCGCGCAGG-CTGTTGGTCCTT--AGGTCGGTGGTTCA--AGTCCGCCCTCACCGGT----
dd_Smed_g4_ASN-GTT_tRNA_5      -GCCT-CCGTGGCGCAATTGGTTAGCGCGTT-CGGCTGTTAACCGAA--AGGTTGGTG--TTC---AGCC-AC-CCGGGCGATT--
dd_Smed_g4_ASN-GTT_tRNA_7      ---GCCGTGAT-CGCTAGTGTTTAGGACATTGCGTTGTTG-CCGCA--ATAACCCAGGTTCG---AATCCTGGTCACGGCA----
dd_Smed_g4_ASN-GTT_tRNA_8      --TGACGTGTAGCTCAGTTGGT---GAGCGGAGGACTGTTAGATCCT-TAGGTCG-TGGTTCA--AATCCGCC-TCACCGCGTGAA
dd_Smed_g4_ASN-GTT_tRNA_9      -CCGGTG-ATAGCTCAGTTGGTTAGCAGCGGAGGACTGTTGATCCTT--AGGTCGGTGGTTCAC-AATCCGCC-TCACCGGA----
                               TSCGKCGYGTAGCKCAGTTGGTTAGGGSSKTAGGRCTGTTRRYCCMAGTAGGTCGGTGGTTCGYGAATCCGMCGTCACGGGADGMA
```

### Formatted Alignments

```
                                   10        20        30        40        50        60        70        80        90
dd_Smed_g4_ASP-GTC_tRNA_2  ATCCTCG--TAG-TATA-GTGGTCAGTA-TCTCC-GCCTGTCACGGCGGGCGACC--CGGGTT-CGATTCCCGGCGCTGATG-
dd_Smed_g4_ASP-GTC_tRNA_3  -TCCTCGAGTAG-TAATTGTGGTCCATGGTCATCCGCCTGTCACGGTGGAAGGCC--CGGGTT-CGATTTCCGGTCGGGGAG-
dd_Smed_g4_ASP-GTC_tRNA_4  -TCCTCGA-TAG-TAA--GTGGTGCATA-TCCTCCGCCTGTCAACGGGAAGGCCC--CGGGTTTCGATTCCCGGTCGGGGAG-
dd_Smed_g4_ASP-GTC_tRNA_5  --CCGTGGATAGCTGTTGGTAGAGCGGA--GGACTGTCAGTTCCTTAGGTCGGCT----GGTTCAAATCCGCCTCATCGGA--
dd_Smed_g4_ASP-GTC_tRNA_6  -TCCTCGA-TAG-TATA-GTGGTCATTA-TCTCC-GCCTGTCACGTTGGAAGGCC---GGGTTCGAATTCCCGGTTCGGGGAG
dd_Smed_g4_ASP-GTC_tRNA_7  -TCCCGTA-TAG-TATA-GTGGTCAGTA-CATCCGCCCTGTCACGCAGGAAAGGCACCGGGTT-CGATTCCCGTCGGGAG---
dd_Smed_g4_ASP-GTC_tRNA_8  -TCCTCGA-TAG-TATA-GTGGTCAGTA-TCTCCCGCCTGTCACG-TGGAAGGCC--CGGGTT-CGATTCCCGGTGGGGAG--
                           ATCCTCGARTAGCTATAKGTGGTCAGTAGTCTCCCGCCTGTCACGKWGGAAGGCCACCGGGTTCCGATTCCCGGTSGGGGRGG
```

### Formatted Alignments

```
                                      10                  20                  30                  40                  50                  60                  70                  80                  90
dd_Smed_g4_CYS-ACA_tRNA_1   - - - - G G T T C G A T G G T G T A G T G T A - - - - - - A T C A C G T - A T G C T T A C A C T A C G C - A G A A G - T T C C C G G T T C G A T C C C G G G T C G A A C G C -
dd_Smed_g4_CYS-ACA_tRNA_2   - - - - G G T T C G A T G G T G T A G C G G T - - - - - - A T C A C G T - C T - C T T A C A C - - - G C - A G A A G G T C C C C G G T T C G A T C C A G G G T C G A A C C A -
dd_Smed_g4_CYS-ACA_tRNA_3   - - G G G G A A C T A G C T - C A C A T G G T - - - - - - A G A G C G C - T C G C T T A C A - - A T G C - - G A G G G T A C T G G G A T C G A T A C C C A G G T T C T C C A -
dd_Smed_g4_CYS-GCA_tRNA_1   - - - - G G G G A T A G C T - C A G - T G G T - - - - - - A G A G C A T - T C G A C T G C A - G A T C G A C T G A G G T C A C G G - T T C A A A T C C G - - - T G C C C C C T
dd_Smed_g4_CYS-GCA_tRNA_10  - - - G G G G T A T T A G C T T C A G T G G T T T - - - - A G A G C A T - T C G A T G G C A - G A T C G - - A G A G G T C C C G G - T T C A A A T C C G A G G T G C C C C T -
dd_Smed_g4_CYS-GCA_tRNA_13  - - T G G G G T A T A G C T T C A T T T G G T - - - - - - A G A G C A T - T C G A C T G C A - G A T C G - C A G A A G T C C C G G - T T C A A A T C C G - G G T G C C C C C T
dd_Smed_g4_CYS-GCA_tRNA_15  - G C C G C G A T A A G C T - C A G T T G G - - - - - - - A G T G C G T - T A G A C T G C A A G A T C T - - A A A G G T C A C T G G T T C G A T A C C G G T T C G C G G C A -
dd_Smed_g4_CYS-GCA_tRNA_2   T A C C G C G T - - A G C T - C A G T T G G G - - - - - - A G A G C G T - T A G A C T G C A - G A T C T - - A A G G T C C A C T G G T T C G A T C C C G G T - C G C G C A A -
dd_Smed_g4_CYS-GCA_tRNA_3   - - C C G G T G A T A G C T - C A G T T G G T - - - - - - A G A G C G G - A G G T T A G C A G T A T C C - - T T A G G T C G G T G G T T C A A A T C C G C C T C A C C G G G -
dd_Smed_g4_CYS-GCA_tRNA_4   - - G G G G G T A T A G - - C T C A G T G G T - - - - - - A G A G C A T - T C T A C T G C A A G A T C G - - A G A G G T C C C C G G T T C A A A T C C G - G G T G C C C C T T
dd_Smed_g4_CYS-GCA_tRNA_5   - - A G G G G T A T G - - - - T C A G T G G T - - - - - - A G A G C A T C T C G A C T G C A - G C T C G - - A G A G G T C C C G G C T T C C A A T C C G - G G T G C C C C T T
dd_Smed_g4_CYS-GCA_tRNA_6   - - G G G G G A A T A G C - - T C A G T G G T A T T A G T A G A G C A T - T C G A C T G C A - G A T C A G A G A G G T C C C C G G T T C A A A A T T C C G G G T G C C C C C C
dd_Smed_g4_CYS-GCA_tRNA_8   - - G G G C G T A T A G C - - T C A G T G G T - - - - - - A G A G C A T - T C G A C T G C A - G A T C T - - A G A G G T C C C C G G T T C A A A T C C G - G G T G C T T C G T
dd_Smed_g4_CYS-GCA_tRNA_9   - - G G G G G T A T A G C T - C A G G T G G T G T - - - - A G A G C A T - T C G A C T G C A - G A T C G A G A G G T C C C C G G G - T T C A A A T C C G - G - T G A C C C C T
                            T R S G G G G T A T A G C T T Y M R G T G G T D T T A G T A G A G C A T C T C G A C T G C A M G A T C S A M A R A G G T C C C S G G T T C A A A T C C G G G G T G C C C C M T
```

### Formatted Alignments

```
                                       10        20        30        40        50        60        70        80        90       100
dd_Smed_g4_GLN-CTG_tRNA_1  GGTCCCATGGT------GTAGCGGTTAG--CACTCTG-ACTCTGAATACAGCGACGCCGACGTTTCAAATCT--CGG-TGGGACCT------------
dd_Smed_g4_GLN-CTG_tRNA_2  ---GGGGTAATAG--CTTCAGT-GGTAGA--GCATTCGACCCTGCAGATCGAGAGGTCCG-GTTCAAA-TCCGGGGTGCCCCT---------------
dd_Smed_g4_GLN-TTG_tRNA_1  ---CCGGTGATAG----GTCAGGGTGAG-----CGAGGACTTTGATTCCTTAGG--TCGGTGGTTCAAATCCGCCTCACCGGA---------------
dd_Smed_g4_GLN-TTG_tRNA_2  AGGTCCATGGTATTTCTGTAGCGGTTAG--CA-TCAGGACTTTGAATCCTGCGA-CCCGA-GGTTCAAATCT--CGG--TGGACTTA-----------
dd_Smed_g4_GLN-TTG_tRNA_3  ---GTCACGATGG---CCGAGTGGTTAAG--GCGTTGGACTTTGAATCCCAAAAAATTGGGGTTTCCCGCGT--AGGTTCGAAACCCTGCTCGTGACG
dd_Smed_g4_GLN-TTG_tRNA_4  ---GGCCCTATAG--TCTCAGG-GGTAGA--GCACTGGTCTTTGTAACCAGGGGGTCGAG-AGTTCGATCTCTCTGGGGCA-----------------
dd_Smed_g4_GLN-TTG_tRNA_5  ---GCCCGGATAG--GCTCAGTCGGTAGA--GCATCAGACTTTGAA--TCTGATGTCCAG-GGTTCGAGTTCCCTGTTCGGGCG--------------
dd_Smed_g4_GLN-TTG_tRNA_6  ---GGGGATGTAG--CT-CAGT-GGTAGA--GCGTCCG-CTTTGCA--TGAAAGGTCCCG-GGTTCGA-TCCCCGGCATCTCCA--------------
dd_Smed_g4_GLN-TTG_tRNA_7  AGTCCCATGGT------GTAGCGGTTAGCACTTCAAGGACTTTGAATCCTGCGACCTCGA-GTTTCAAATCT--CGG-TGGGACCT------------
dd_Smed_g4_GLN-TTG_tRNA_8  GGTCCCATGGTT---CTGTAGCGGTTTAGCCACTCAGGACTTTGAATCCTGCGA-ACCGA-GTTCAAATCTT--CGGCTGGGACCT------------
                           RGTSCCRTGRTAGTTCTGYAGYGGTTAGAMCGCWYWGGACTTTGAATCCYGMGAG CCGGBGKTTCAADTCYSCBGGDTSGGACCTMTGCTCGTGACG
```

### Formatted Alignments

```
                          10        20        30        40        50        60        70        80
dd_Smed_g4_GLU-CTC_tRNA_2 TCCCTGATGGTC-TAGCGGTTAGGATTCCTGGCTCTCACCCAGGTGGCCC-GGGTTCGACTCCCGGTCAGGAAT
dd_Smed_g4_GLU-CTC_tRNA_3 TTCCTGATGGTC-TAACGGTTAGGATTCCTGGCTCTCACCCGGGTAGCCT-GGGTTCGACTTTCGGTCAGGAAA
dd_Smed_g4_GLU-TTC_tRNA_1 TCCCTGATGGTCCTAGCGGTTAGGATTCCTGGTTTTCACCCAGGCGGCCCCGGGTTCGACTCCCGGTCAGGGAT
                          TCCCTGATGGTCCTAGCGGTTAGGATTCCTGGCTCTCACCCAGGTGGCCCCGGGTTCGACTCCCGGTCAGGAAT
```

### Formatted Alignments

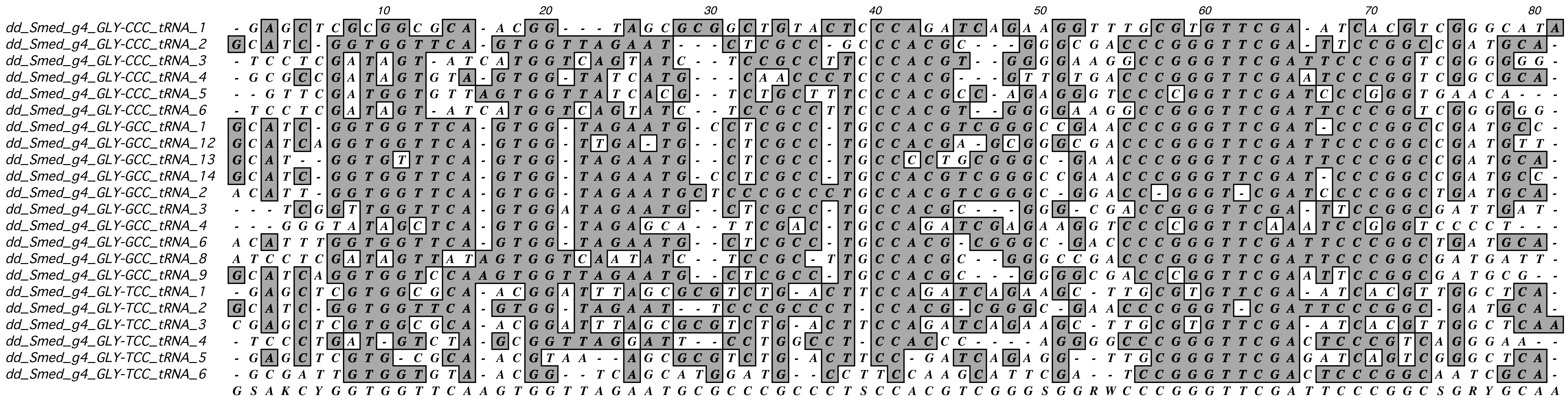

### Formatted Alignments

```
                                    10        20        30        40        50        60        70        80        90        100
dd_Smed_g4_HIS-GTG_tRNA_1  --GTGCGTG-TCG-TCTA-GTGGT-AGGACATTGCGTTGTGGCCGCAATA-----CCAGGTTTCGAATCT--GTCACGGCACT
dd_Smed_g4_HIS-GTG_tRNA_2  ---GCCGTGATCGTTCTA-GTGGTTAGGACATTGCGTTGTGGCCGCAATAA----CCCAGGTTCGATCCAT-GG--CGGCA--
dd_Smed_g4_HIS-GTG_tRNA_3  --GCCCGTGATCGGTCTAAGTGGTTAGGAACTTGCGTTGTGGCCGCAATAA----CCCAGGTTCGAATCCT-GGTCCGGCA--
dd_Smed_g4_HIS-GTG_tRNA_6  GTGGCCGTGATCGCAGTG-GTAA--GGGACATTGCGT-GTGGC-GCAATAA----CCCAGGTTCGAATCCT-GGCACG-CACT
dd_Smed_g4_HIS-GTG_tRNA_7  ---GCCGTGATCG-TCTA-GTGGTTAGGACATTGCGTTGTGGCCGCAAGGAGGAGCCCAGGTTCGAATCCTGGTCAAGGCA--
                           GTGGCCGTGATCGBTCTAAGTGGTTAGGACATTGCGTTGTGGCCGCAATAAGGAGCCCAGGTTCGAATCCTGGGCACGGCACT
```

### Formatted Alignments

```
                                             10                  20                  30                  40                  50                  60                  70                  80
dd_Smed_g4_ILE-AAT_tRNA_1   - G G C C G A G T A G C T C A G T T G G T T A G A G C G T C G T G C T A A T A - A C G C G A A T G T C G T G G G T T C G A G C C C C A T C T T G G C C A -
dd_Smed_g4_ILE-AAT_tRNA_4   - G C C T T C G T A G C T C A G T - G G T T A G A G C A C A G G T C T A A T A - A A C C A A T G G T C G T G A G T T C A A T T C T C A C C G A A G G C A -
dd_Smed_g4_ILE-TAT_tRNA_1   - T G C C C C G T A G C T C A G T C G G T C A G A G C G C T G T A C T T A T A - A T G C G G A G G T C G A G A G T T C A A G C C T C T C T C G G G G C A C
dd_Smed_g4_ILE-TAT_tRNA_2   G G T T C G A T T G G T G T A G T - G G T T A T C A C G T C T G C T T T A T A C G C A G A A A G G T C C C C G G T T C G A T C C C G G G T C G A A C C A -
                            G G B C C B M G T A G C T C A G T Y G G T T A G A G C G Y H G K   C T W A T A C A H V C R A A G G T C G H G R G T T C R A K C C Y C D B Y B D R G S C A C
```

### Formatted Alignments

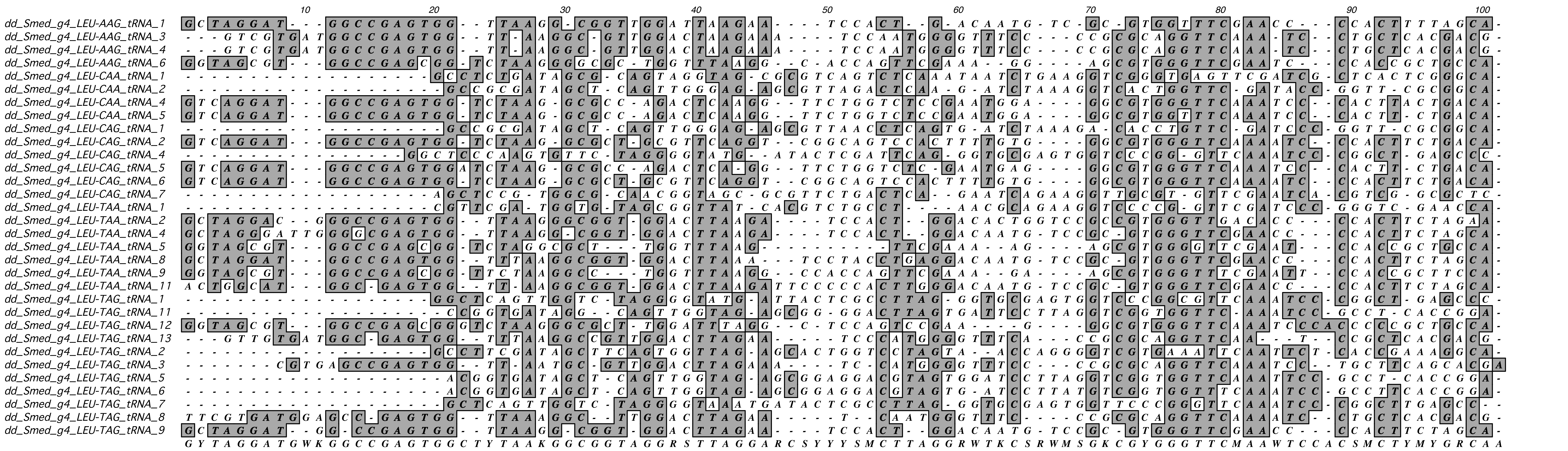

#### Formatted Alignments

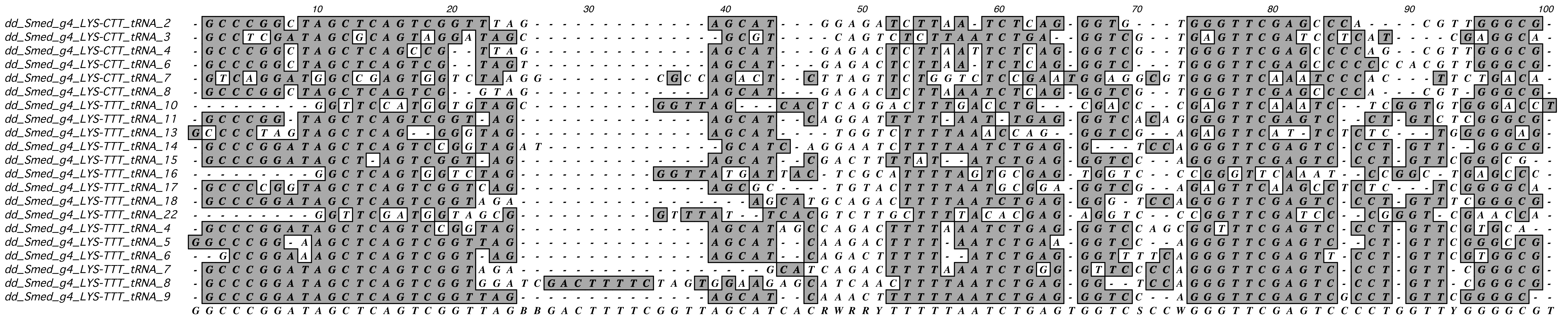

### Formatted Alignments

```
                                       10        20        30        40        50        60        70        80        90       100
dd_Smed_g4_MET-CAT_tRNA_2    -----TGATTAGCGCAGTAGGTTAGC-----GCGT-C-AGTCTCATAA-TCTGCA-AGGTTCGTGAGTTCGATTCTCATGGGCATAA
dd_Smed_g4_MET-CAT_tRNA_3    --GTCAGGATGGCCGAGTGGT-TAAGGCGCCAGACTCATTTTTCTGGTCTCCGAATGGAGGCGTGGGTTCAAATCCCACTTCTGACA
dd_Smed_g4_MET-CAT_tRNA_5    -GCCTCGATTAGCGCAAGTAGGTAGC-----GCGT-C-AGTCTCATAA-TCTGCA-GGTTTCGTGAGTTCGATCCTCACTCGGGGCA
dd_Smed_g4_MET-CAT_tRNA_6    AGCCTCG--TAGCGCAGTTAGGTAGC-----GCGT-CCAGATTCATAA-TCTGTA-AGGT-CGTGAGTTCGATCCTCACTCGGGCAA
dd_Smed_g4_MET-CAT_tRNA_7    -GCCTCGA-TAGCGCAGTAAGGTAG------GCGT-CAGTCCTCATAA-GTCTGA-AGGT-CGTGAGTTCGATCCTCATTCGAGGCA
dd_Smed_g4_MET-CAT_tRNA_8    --GTCAGGATGGCCGAGTGGTCTAAGGCGCCAGACTCATGTT-CTGGTCTCCGAATGGAGGCGTGGGTTCAAATCCCACTTCTGACA
dd_Smed_g4_iMET-CAT_tRNA_1   -AGCAGA-GTGGCGCAGTGGAAA--------GCGTGCTGGGCCCATAAACCCAAG-AGGTCCGTGGATCGAAACCACGCT-CTGCTA
dd_Smed_g4_iMET-CAT_tRNA_2   --GCAGAAGTGGCGCAGTGGAAGCGTGCTGGGCCTGCTGGGCCCATAA-CCCAGA-GGTTCCGTGGATCGAAACCACGCTTCTGCT-
dd_Smed_g4_iMET-CAT_tRNA_3   -AGCAGA-GTGGCGCAGTGGAAA--------GCGTGCTGGGCCCATAAACCCAAG-AGGTCCGTGGATCGAAACCACGCT-CTGCTA
                             AGGCWSGADTGGCGCAGTGGRRTAGSGCGCCGCGTGCWRGKCTCATAAMTCCGVATAGGTBCGTGGGTTCAAACCWCACTYCTGVYA
```

### Formatted Alignments

```
dd_Smed_g4_PHE-AAA_tRNA_1  --GGTAGCGTGGCCGAGCGTGTCTAAGGCGCTGGTTTAAAGGTTCGAAAGAA--------GCGTGGGTTCG---AATCCC--ACCGCTGCCA
dd_Smed_g4_PHE-AAA_tRNA_2  --GGTTCGATGGTGTAGCGGTTAT-GCACGTC-TGCCTAAAACGCAGAAGGT---------CCCCGGTTTC---GATCCG--GGTCGAACCA
dd_Smed_g4_PHE-AAA_tRNA_3  --GTCGTGATGGCCGAG--TGGTTAAGGCGTTGACTAAAAAATTCCAATGGG---TTTCCGCGCAGGTTTTTCAAATCCT--GCTC-CGACG
dd_Smed_g4_PHE-AAA_tRNA_4  --GCCGCGATAGCTCA-TTGGGA--GAGCGTTAGACGAAAGATCTACAAGGT---------CACTGTTC-G---ATTCCC-GGTTCGCGGCA
dd_Smed_g4_PHE-AAA_tRNA_5  --GTCGTGATGGCCGAG--TGGTTAAGGCGTTGGACTAAAATT-----------TTCACCGCGCAGGTTCA--AATGCCT--GCTCACGACG
dd_Smed_g4_PHE-AAA_tRNA_6  --GGTTCGATGGTGTAGCGGTTATCAAACGTCCTGCCTAAACCGCAGAAGGT---------CCC-GGTTTG---ATCCCG--GGTCGAACCA
dd_Smed_g4_PHE-GAA_tRNA_1  --GCCGCGATAGCTCAGCTGGGA--GAGCGTTAGACTGAAGATTCTAAAGGT---------CGCTGGTTTC---GATCGC--GTTCGCGGCA
dd_Smed_g4_PHE-GAA_tRNA_3  --GCCGCGATAGCTCAGTTGGGA--GAGCGT--GACTGAAGAATCTAAAAGG---------CACTGGTT-C---GATCCCTGGTTCGCGGCA
dd_Smed_g4_PHE-GAA_tRNA_5  --GTCGTGATGGCCGAG--TGGTTAAGGCGTTGGACAGAAAAT-CCAATGGG--TTTCCGCGCAGGTTC----AATCCT--GCTCACGACG
dd_Smed_g4_PHE-GAA_tRNA_6  --GTCACGATGGCCGAG--TGGT-AAGGCGTTGGTT--GAAAT-CCAATGGGGTTTCCCCGCGTAGGTTC----G--CCT--GCTCGTGACG
dd_Smed_g4_PHE-GAA_tRNA_7  GTTTATCGATAGCTCAGTTGGGA--GAGCGTTAGACTGAAGATCTA-AAGGT---------CACTGGTT-C---GATCCC-GGTTCGCGGCA
                           GTGBCGCGATGGCYSAGCTGGGATAARGCGTTGGACHRAAAATBCMAAAGGKGTTTYHCCGCGCWGGTTYSTCAAATCCYTGGYTCGCGRCA
```

### Formatted Alignments

```
                                    10        20        30        40        50        60        70        80        90       100
dd_Smed_g4_PRO-AGG_tRNA_1  GGCTCAGTG-GTCTA-GGGGTATGATACCTC-GCTTAGGGTGCGAG---TGGT------CCCGGGTTCCAAATCCCGGC-TGAGCCC
dd_Smed_g4_PRO-AGG_tRNA_2  GGTAGCGTGGCCGAGCGGTCTAAGGCGCTGG-TTTAGGCGTTCGAA---AGA-------GCGTGGGTTCGAATCCCACCGCTGCCA-
dd_Smed_g4_PRO-AGG_tRNA_5  GTCGTGATT-GCCGA-TGTTTAAGGCGTTGGATCTAGGAAATCCAT---GGGTTTCCCCGCGCAGGTTCAAATCTTGCT-CACGACG
dd_Smed_g4_PRO-CGG_tRNA_1  GGCTCAGTGAGTCTAGGGGTTATGATTCTCG-CTTCGGGTGGCGAG---AGGT------TCCCGGGTTCAATTCCCGGC-TGAGCCC
dd_Smed_g4_PRO-GGG_tRNA_1  GGCTCAGTG-GTCTA-GGGGTATGATTCTCG-CTTGGGGGTGCGAG---AG---------TCCCGGTTCAAAATCCGGC-TGAGCCA
dd_Smed_g4_PRO-TGG_tRNA_1  GTCACGATG-GCCGAGTGGTTAAGGCGTTTGACTTGGAAATCCAATGGAGGGTTTCCCCGCGTAGGTTCGAACCCTGCTTCGTGACG
dd_Smed_g4_PRO-TGG_tRNA_2  GTCACGATG-GCCGAGTGGTTAAGGCGTTTGACTTGGAAATCCAATGGAGGGTTTCCCCGCGTAGGTTCGAACCCTGCTTCGTGACG
dd_Smed_g4_PRO-TGG_tRNA_3  GGCTCAGTG-GTCTAGGGGTAATGATTCTCG-CTTTGGGTGGCGAG---AG--------TCCCGGGTTCAAATCCCGGC-TGAACAG
dd_Smed_g4_PRO-TGG_tRNA_4  GCCGTGATCGATCTA-GTGGTAGGACATGCG-TGT-GGTCCGCAAT---AA---------CCAAGGTTCGAATTCCTGGTCACGGCA
dd_Smed_g4_PRO-TGG_tRNA_6  GGCTCAGTG-GTCTAGGGGTAATGATTCTCG-CTTTGGGTGGCGAG---AG--------TCCCGGGTTCAAATCCCGGC-TGAACAG
dd_Smed_g4_PRO-TGG_tRNA_7  -GCTCAAGTGGTCTACGGGGTATGAT-CTCG-CTTTGGGT-GCGAG---AGG-------TCCCGGGTTCAAATCCCGGCCTGAGTC-
dd_Smed_g4_PRO-TGG_tRNA_8  GGCTCAGTG-GTCTAAGGGGTATGATTCTCG-CTTTGGGGTGCGAG---AGG--------TCCGGGTTCAAAATCCCGC-TAGCCC-
                           GGCTCAGTGGGTCTAGGGGKTATGATKCTCGACTTKGGGKTGCGAGGGAAGGTTTCCCCKCCCGGGTTCAAATCCCGGCTTG GCCG
```

### Formatted Alignments

```
                          10        20        30        40        50        60        70        80        90        100
dd_Smed_g4_SEC-TCA_tRNA_1 GCGGGCATGAGCCTCGGCGGTCCGGGGTGCAGGCTTCAAACCTGTAGTCGGTTGACACCGAAGTGGTTCGATTCCAACTTTCCAGC-
dd_Smed_g4_SEC-TCA_tRNA_2 GCTGGGATGAGCCTCGGCGGTGCGGGGTGCAGGCTTCAAACCTGTAGTTGGTTGACACCGAAGTGGTTCGATTCCACCTTTCCAGCG
                          GCKGGSATGAGCCTCGGCGGTSCGGGGTGCAGGCTTCAAACCTGTAGTYGGTTGACACCGAAGTGGTTCGATTCCAMCTTTCCAGCG
```

### Formatted Alignments

```
                                   10        20        30        40        50        60        70        80        90       100
dd_Smed_g4_SER-ACT_tRNA_1  -GACGA-GGTGGCCGAGTGG-TTAA-GGCGATGGACTACTAATCC-ATTGGGGTTTCCCC---GCGTGGGTTCGAATC--CCATCCTCGTCG--
dd_Smed_g4_SER-ACT_tRNA_2  --GCCTTCGTAG-CTAGTGG-TTAG-AGCACTGGTCTACTAAACC---AGGGGTCGT----------GAGTTCAATTC--TCACCGAAGGCA--
dd_Smed_g4_SER-ACT_tRNA_3  ----------GTTCGAATGGTGTAA-----GCGGTTATCACGTC---TGTTTACTACGCAGAAGTTCCCGGTTCGATC--CCGGGTCGAACG--
dd_Smed_g4_SER-AGA_tRNA_1  -ATCGT-GATGGCCGA-GTGGTTAA-GGCGTTGGACTAGAAATCC-AATGGGTTTCCCC--GCGCAGTTGTTCAAATC--CTGCTCACGCACG-
dd_Smed_g4_SER-AGA_tRNA_2  --TCGT-GATGGGCGAGTGGTTTAA-GGCGTTGGACTAGAAATCCAATGGGGGTTTCCC--CGGCAGG--TTCAAATC--CTGACTCACAAAG-
dd_Smed_g4_SER-AGA_tRNA_3  -GTCGT-GATGGCCGAAGTGGTTAG--GCGTTGGACTAGAAATCCAAATGGCGGTTTCCCCGCGTAGG-TCTCAAATC--CTGCTCACGAGC--
dd_Smed_g4_SER-AGA_tRNA_4  ----GTAGATGGCCGAGTG-TTTAA-GGCGTTGGACTAGAAATCCGA-ATGGGTTCCCC--GCGCAGG--T-CAAATC--CGGCTCAACG----
dd_Smed_g4_SER-AGA_tRNA_7  CGTGAT-ATGCGCCGAGTGTTATAG-TGCGTTGGACTAGAAATCC-AATGGGGTTTCCCCTGCGCAAG-GTTCAA-AT--CCTGTCACGA----
dd_Smed_g4_SER-AGA_tRNA_8  --GGTAGCGTGGCCAAGCGGTCTAACGGCGCTGGTTTAGAGGCTCCA--------GTCCGAAGGCGTGGGTTCAAATC--CCACCGCTGCCA--
dd_Smed_g4_SER-AGA_tRNA_9  -GTCGT-G-ATGCCGAGTGGTTTAG-GGT-TGGGACTAGAAATC--AATGGGGTTTCCC----GCAGG-TTTCAA-TC--CTGCTCACGACG--
dd_Smed_g4_SER-CGA_tRNA_2  CGTCACG-ATGGCCGAGTGG-TTAA-GGCGTTGGACTCGAAATCC-AAGGGGTTTTTCCC-CGCGA-GAGTTCAAATC--CTGCTGTGACGA--
dd_Smed_g4_SER-CGA_tRNA_4  -GTCACG-ATGGCCGAGTTG-TCAA-GGCGTTGGACTCGAAATCC-AATGTGGTTTCCCC-GCGC--AGGTTCAAATC--CTGC--TCGTGACG
dd_Smed_g4_SER-CGA_tRNA_5  --------GGTGCCGAGTGGTATAG--GCGTTGGACTCGAAATCC-AATGGGGTTTCCCC-GCGCAGG-TTCAAA-TC--CTGCTCT-------
dd_Smed_g4_SER-CGA_tRNA_7  --TCACGGATGGACGAGTGG-TTAA-GGCGTTGGACTCGAAATCC-AATGGGTTTCCCGC-GC----AGGTTCAAAATACCTGC--TCGTGAA-
dd_Smed_g4_SER-GCT_tRNA_1  -GACGA-GGTGGCCGAGTGG-TTAA-GGCGATGGACTGCTAATCC-ATTGGGGTTTCCCC---GCGTGGGTTCGAATC--CCATCCTCGTCG--
dd_Smed_g4_SER-GCT_tRNA_2  -CCGGTGATAAGCTCAGTTGGTAAT-AGCGCGAGGACTGCTACCT-CTA--GGTCGTGTG---------GTTCAAATC--CGCCTCACCGGA--
dd_Smed_g4_SER-TGA_tRNA_2  -GCCGCGATA-GCTCAGTTGGAGA--AGCGTTAGACCTGAAAGAT-CTAAAGGTCAC-TG---------GTTCGATCC--CGGTTCGCGGCA--
dd_Smed_g4_SER-TGA_tRNA_3  ----------GGTCCATTGGTGTA------GCGGTTTACACTCTCAGGACTTGAAATCCTGCGACCCGAGTTCAAATC--TCGGTGGGACCA--
dd_Smed_g4_SER-TGA_tRNA_4  -TCCGT-GATGGCCGAAGTGGTTAA-GGCGTTGGACTG--AATCC-AATGGGGTTCCC---GCGCAG--GTTCAAATC--CTGCTCACCGAC--
dd_Smed_g4_SER-TGA_tRNA_5  -GCCGCGATA-GCTCAGTTGGAGA--AGCGTTAGACCTGAAAGAT-CTAAAGGTCAC-TG---------GTTCGATCC--CGGTTCGCGGCA--
dd_Smed_g4_SER-TGA_tRNA_6  --TCACGCATGC-CGAGTGG-TTAA-GGCGTTGGACTTGAAATCC-AATGGGGTTTCCCC-GCGTT-AGGTTCGGAAC-CCTGCCTTCGTGAA-
dd_Smed_g4_SER-TGA_tRNA_7  -GTCACG-ATGGCCGAGTGG-TTAA-GGCGTTGGACTTGAAATCCCAATGGGGTTTCCCC-GCGTTTAGGTTCGAACC--CTGCTCGTTTGACG
dd_Smed_g4_SER-TGA_tRNA_9  ----------GGTCCCATGGTGTA------TGCGTTAGCACTCT-GGAATCTGAAATCCAGCGACC-GAGTTCAAATC--TCGGTGGGAACT--
                           CGTCGYGGATGGCCGAGTGGTTTAACGGCGTTGGACTWGAAATCCVAATGGGGTTWCCCC GCGCAKGGGTTCAAATCACCYGCTCWCGDSAVG
```

### Formatted Alignments

```
                                    10        20        30        40        50        60        70        80        90       100
dd_Smed_g4_SUP-CTA_tRNA_1   ----CTTCGTTAGCTCAGT--GGTT--AGAGCACTGGTCTCTAGTAAACCAGGGGT--------CGT----GAGTTCAA-TTC---TCACCGAAGC
dd_Smed_g4_SUP-CTA_tRNA_3   ---GCCTCGATAGCGCAGT-AGGT----AGCGCTCAGTCTCTAAATCTGAAGGCGT--------------GAGTTCGATCCTCACTCGGGGCA---
dd_Smed_g4_SUP-CTA_tRNA_4   ---GGTTCGATGGTGTAAG-TGGTT--AATCACGTCGCTTCTAGACGCAGAAGGTC-------------CC-GGTTCGATCCCGGGTCGAACCA--
dd_Smed_g4_SUP-TCA_tRNA_1   ---TCCTCGATAGTA-TAG-TGGTC--AATAATCTCCGCCTTCAC---CGTGAAGGC-------------CGGGTTCGATTCCCGTCGGGGAG----
dd_Smed_g4_SUP-TCA_tRNA_2   ---TCCTCGATAGTA-TAG-TGGTC--AATAATCTCCGCCTTCAC---CGTGAAGGC-------------CGGGTTCGATTCCCGTCGGGGAG----
dd_Smed_g4_SUP-TCA_tRNA_3   ---GGTTCGATGGTTGTAG-TGGTT--TATCA-ACGTCTGCTTCACGCAGAAGTTC-------------CCCGGTTCGATCCCGGGTTCGACCA--
dd_Smed_g4_SUP-TTA_tRNA_10  ---GCTAGGATGGCCGAA--TGGTGGTAAGGCGGTGGACTTTAAAATCCACTGGACAATTGTCTCGC-GTGGGTTCGAA-CCCCACTTTCTAGCA-
dd_Smed_g4_SUP-TTA_tRNA_11  -CCGGTGTGATAGCTCAGTTTGGTA--CGAGCGGAGGACGTTAGTATCCTTAGGTC--------GGT----GGTTCAAA-TCCGCCTCACCGGG--
dd_Smed_g4_SUP-TTA_tRNA_18  ---GGTCCCATGGTG-TAG-CGGT---AGCACT-CAGGACTTTAAATCCTGCGACCC--------CT--CGAGTTCAAATCTCGGTGGGACCT---
dd_Smed_g4_SUP-TTA_tRNA_19  ---GTCGTGATTGCCGAAG-TGGTT--AAGGCGTTG-ACTTTAAAATCCAATGGGT---TTCCCCGC-GCAGGTTCAAAATCCTGCTCACGACG--
dd_Smed_g4_SUP-TTA_tRNA_2   ---GGTTCGATGGTGTAAG-CGGTT---ATCA--CGTCTGCTTTAACGAGAAGGTC-------------CCCGGTTCGATCCCGGGTCGAACCA--
dd_Smed_g4_SUP-TTA_tRNA_20  ---GGTTCGATGGTGTAAG-CGGTT---ATCA--CGTCTGCTTTAACGAGAAGGTC-------------CCCGGTTCGATCCCGGGTCGAACCA--
dd_Smed_g4_SUP-TTA_tRNA_21  GGCTACACAGTGGTC-TAG-GGGTA--TGATACCTCGCTTTTAGGTGCGAGTGGTC-------------CCGGGTCAAATCCCGGCTGAGCCC---
dd_Smed_g4_SUP-TTA_tRNA_22  ---GTCACGATGGCCGAG--TGGT---AAGGCGTTGGACTTTAAA-TCCAATGGGG---TTTCCCGCCGTAGGTTCGAA-CCCTGCTCGTGACG--
dd_Smed_g4_SUP-TTA_tRNA_4   ----GCCCGGAAGCTCAGG-TCGT----AGAGCAATCAGACTTAATCTGAAGGTCCA-------------GGGTTCGAGTCCTGTTCGGGCG----
dd_Smed_g4_SUP-TTA_tRNA_5   -----CACGACGACCGAG--GGGTT--AAGGCGTTGGACTTTAAA-TCCAATGGGG---TTTCCCGC-GTAAGTTCGAA-CCCTGCTCGTGA----
dd_Smed_g4_SUP-TTA_tRNA_6   ---GGTCCCATGGTG-TAG-CGGTT--AGCACTTCAGGACTTTAATCTCTGCGACC-------------CGAGTTCCAAATCTCGTGGAACCT---
                            GSCGSTWCGATGGTSSAAGTTGGTTGTAARVMSKYGGMCTTTARAAYCSWAWGGKCAATTTTCCCGCCGCSGGTTCSAATCCCGGBTSGRMSCARC
```

### Formatted Alignments

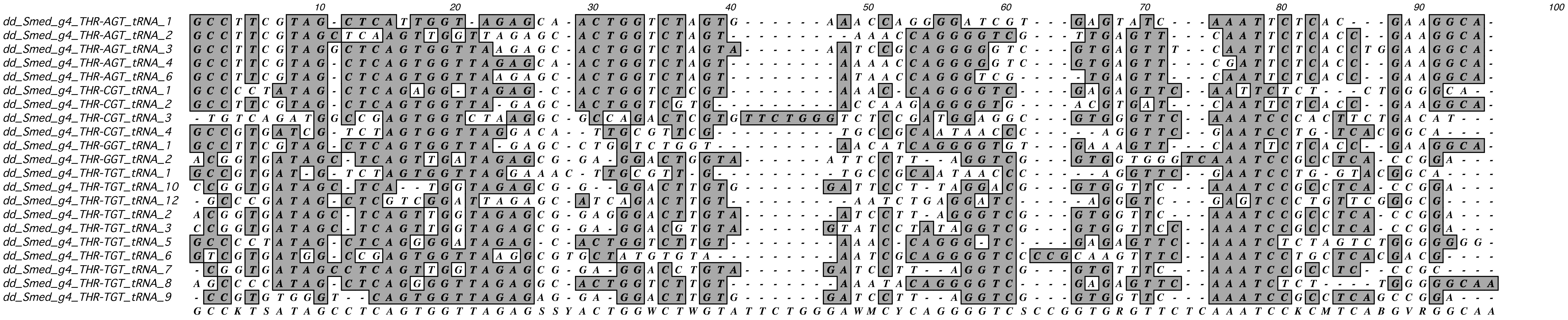

### Formatted Alignments

```
                                  10        20        30        40        50        60        70        80        90        100
dd_Smed_g4_TYR-ATA_tRNA_2  ---GGCTCAGTGG-TCTAGG--GGTAATGATACTCGCTGATAGGGTGCGAAGT---GGTCCCGGGTTCAAAATCCCGCTGAGCCC--
dd_Smed_g4_TYR-ATA_tRNA_3  ---GCCTTCGTAAGCTCAGT--GGTTAGAGCACTGGTCTATAAAAACCAGGGG-----TCGTGGAGTTCAATTCTCACGAAGGCA--
dd_Smed_g4_TYR-GTA_tRNA_1  CCGGTG---ATA--CTCAGTTGG--TAGAGCGGAGGACTGTAATTCCTTAGCG----TCCGAGTGGTTCCAATCC-GCCTCACCGGA
dd_Smed_g4_TYR-GTA_tRNA_2  ---TCCTCGGTAG-TATAGT--GGTCAGTATCTCCGCCTGTACAGCTTGGAA----GGCGCCGGGTT--GATTCCCGGTCGGGGAG-
dd_Smed_g4_TYR-GTA_tRNA_3  --CGTG---ATAC-CTCAGT-GG--TAGAGCTGAGGACTGTAGGATCCTTAGGGTTTTTTCGGTGGTTCAAATCC-GCCTCACACGA
dd_Smed_g4_TYR-GTA_tRNA_4  CGGGTG---ATAGCCTCAGTTCG--TAGAGCGGAGGACTGTATGATCCTTAGG------TCGGTGGTTCAAATCCCGCCTCACCAGG
dd_Smed_g4_TYR-GTA_tRNA_5  CCGGTG--GATAG-CTCAGT--G--TAGAGCGGATGACTGTAGAATCCTT--A----GGTCGGTGGTTCAAATCC-GCCTCACCGGA
dd_Smed_g4_TYR-GTA_tRNA_6  ---GTCTTCGTA-GCTCAGT--GGTTAGAGCACTGGTCAGTAAAA-CCATGGG-----TCGTG-AGTTCAATCTCAACGAAGGCA--
dd_Smed_g4_TYR-GTA_tRNA_7  ACGGTG---ATAG-CTCAGTTGGCTTAGAGCGGAGGACTGTAGGATCCTTAGG----TCCGTTGGTTTCAAATCC-GCCTCACCGGA
dd_Smed_g4_TYR-GTA_tRNA_9  CCGGTGTTGATAG-CTCAGTT-G--GAGAGCGGAGGACTGTAAGATCCTTTAA----AATCGGTGGTTCAAATCC-GCCTCACCGGA
                           CCGGTGTTSATAGGCTCAGTTGGGTTAGAGCRGAGGACTGTARRATCCTTAGGGTTGKYYCKGTGGTTCAAATCCCGCCTCACCRGA
```

### Formatted Alignments

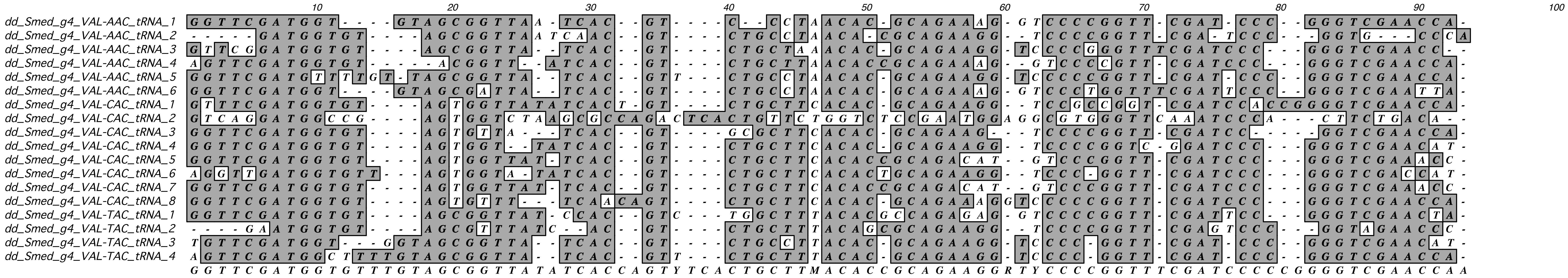

Formatted Alignments

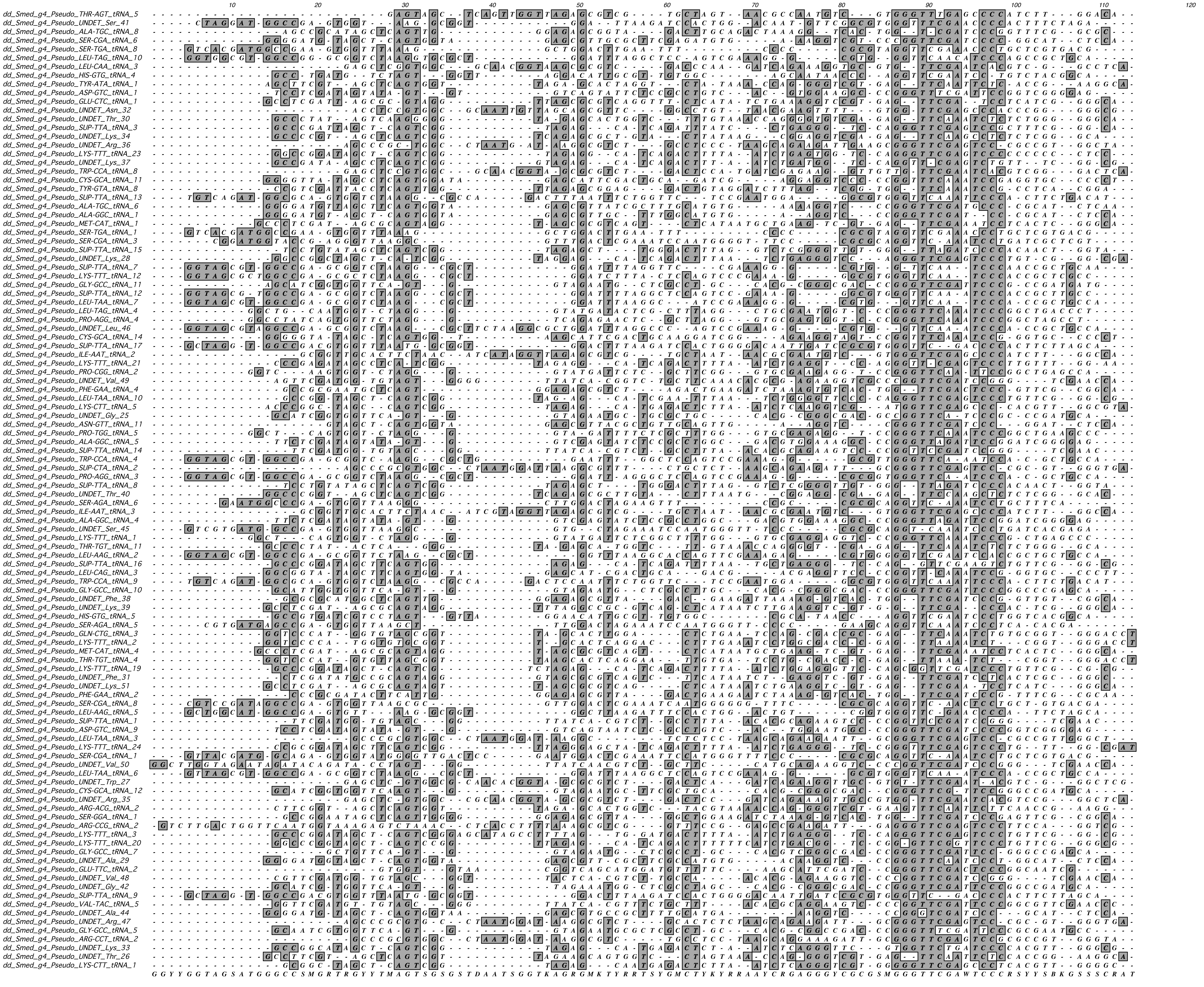

### Formatted Alignments

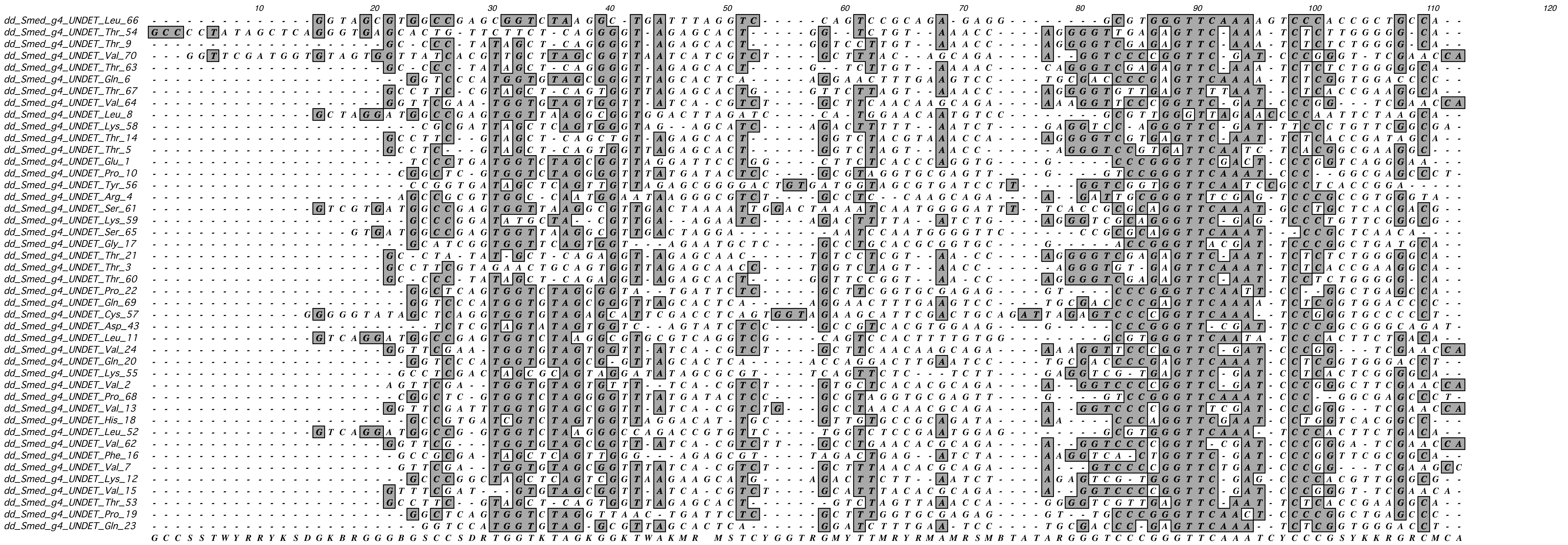
